# Astrocyte targeted SMN1 gene therapy and forskolin application improves astrocyte filopodia actin defects and motor neuron synaptic dysfunction in human SMA disease pathology

**DOI:** 10.64898/2026.03.26.714618

**Authors:** Emily Welby, Xiaojie Liu, Melinda Wojtkiewicz, Linda Berg Luecke, Rebekah. L. Gundry, Qing-Song Liu, Allison. D. Ebert

## Abstract

**Background:** Peri-synaptic astrocyte processes (PAPs) play a fundamental role in synapse formation and function. Central afferent synapse loss and astrocyte dysfunction greatly impede sensory-motor circuitry in spinal muscular atrophy (SMA) disease progression, however mechanisms underpinning tripartite synapse dysfunction remains to be fully elucidated. The aims of this study were to further define PAP and motor neuron synaptic defects in human SMA disease pathology and implement a therapeutic intervention strategy to improve motor neuron function.

**Methods:** We derived astrocyte monocultures and motor neuron astrocyte co-cultures from healthy and SMA patient induced pluripotent stem cell (iPSC) lines to assess intrinsic astrocyte filopodia defects and phenotypes occurring at the synapse-PAP interface, respectively, using cell surface capture mass spectrometry proteomics, confocal and super resolution microscopy, synaptogliosome isolation, and electrophysiology.

**Results:** SMA astrocytes demonstrated intrinsic filopodia actin defects featuring low abundance of actin-associated cell surface N-glycoproteins, and decreased filopodia density and CDC42-GTP levels after actin remodeling stimulation. This phenotype is likely driven by the significant reduction of CD44 and phosphorylated ezrin, radixin and moesin ERM proteins (pERM) within SMA astrocyte filopodia. The dual combination of SMN1 gene therapy and forskolin treatment, an adenylyl cyclase activator leading to increased cyclic adenosine monophosphate (cAMP) levels and actin signaling pathway stimulation, led to extensive branching and increased filopodia density of SMA astrocytes during actin remodeling. SMA patient-derived motor neuron and astrocyte co-cultures, particularly samples derived from male patient iPSC lines, demonstrated a significant decrease in synapse number, actin-associated pre-synaptic neurotransmitter release protein, synapsin I (SYN1), and PAP-associated expression of pERM and glutamate transporter, EAAT1. Our astrocyte-targeted SMN1 augmentation and forskolin treatment paradigm restored SYN1 protein levels within the SMA synaptogliosome, resulting in significant increases in motor neuron synapse formation and function, but did not fully restore PAP-associated proteins levels at the synapse.

**Conclusions:** SMA astrocytes demonstrate intrinsic actin-associated defects within filopodia, which correlates with decreased pERM levels at tripartite motor neuron synapses. We also define a SMN- and cAMP-targeted treatment paradigm that significantly increases pre-synaptic neurotransmitter release protein levels to improved SMA motor neuron synapse formation and function.

**Graphical abstract:** 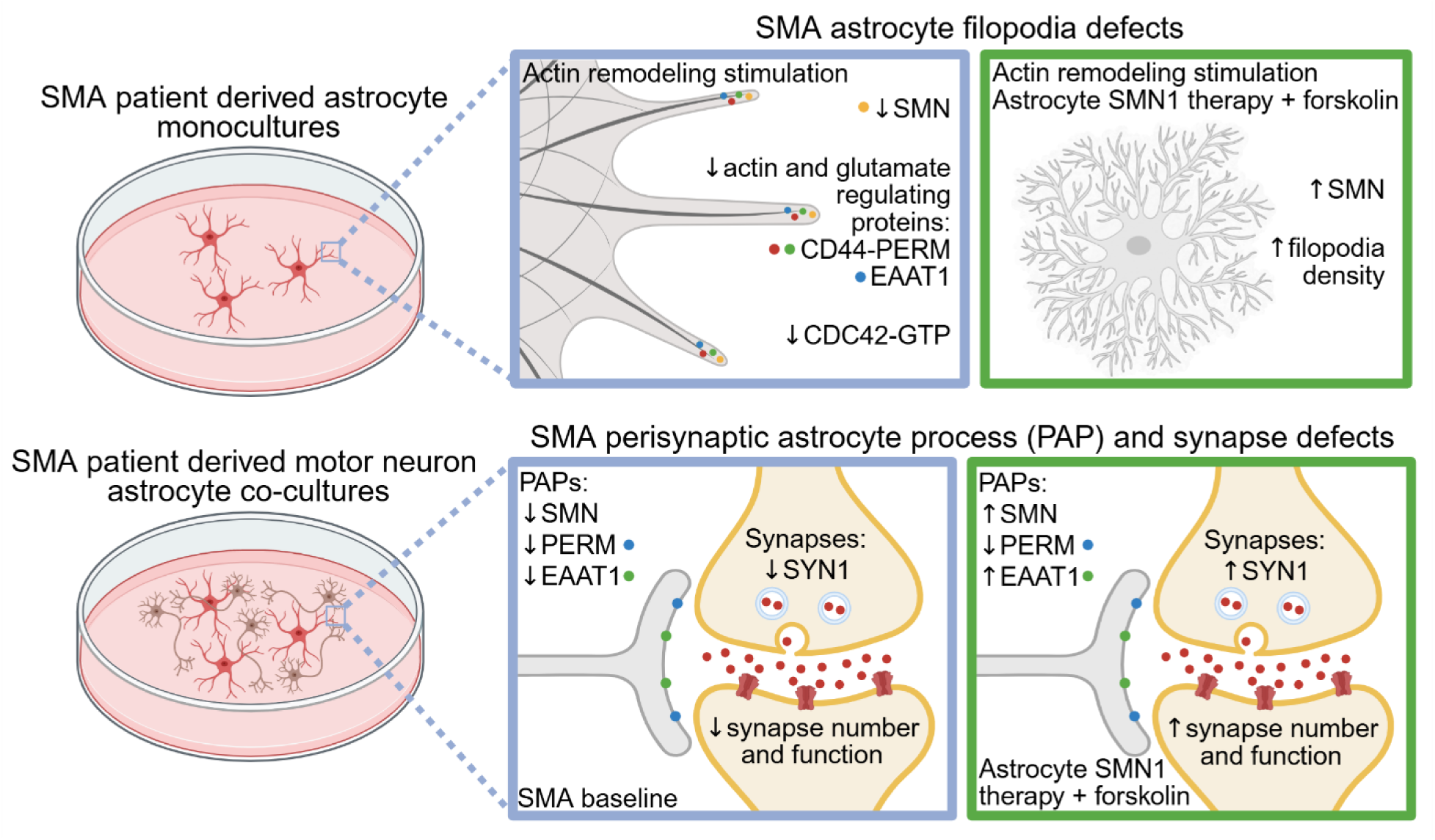

## Background

Astrocytes directly interact with neuronal synapses through peri-synaptic astrocyte processes (PAPs), which are extremely fine filopodia responsible for the dynamic regulation of synaptic structure, development and function (1–6). To perform these roles, PAPs are uniquely enriched with mRNAs that encode cell surface-associated proteins to guide synaptogenesis (i.e., SPARC (7), sense synaptic activity (i.e., metabotropic receptors (8)), and facilitate neurotransmitter and ion clearance from the synaptic cleft (i.e., Na^+^ dependent excitatory amino acid transporter (9)). These allow for a bi-directional communication where neuron-released neurotransmitters evoke glial intracellular calcium transients to elicit the PAP-mediated regulation of synaptic events, such as gliotransmitter release and PAP outgrowth or retraction (10–13). The actin cytoskeleton is solely responsible for PAP remodeling; specifically, the phosphorylated state and protein interactions of Ezrin, a member of the ERM (ezrin, radixin and moesin) protein family, are thought to be crucial in regulating PAP actin dynamics (8, 14). The activated form of ezrin (phosphorylated ezrin threonine-567) is highly enriched within PAPs (8, 14), where it acts as linker to tether actin filaments to the plasma membrane to facilitate the regulation of filopodia formation and motility (15, 16). In other cell types, phosphorylated ezrin regulates actin dynamics through the interaction of cell surface adhesion molecules, e.g., CD44 (17–19), allowing the downstream activation of actin remodeling pathways, such as CDC42 (20–22). In relation to synaptic plasticity, it is hypothesized that ERM protein complexes help stabilize hyaluronic acid-CD44 interaction and actin filaments at the plasma membrane to support synapse formation and neural circuit function via actin dynamics (23).

Vast diversity in synaptic features, such as morphology, molecular composition and contact with PAPs, exists across the CNS that is influenced by anatomical region, age and neural circuitry activity (24). Specifically, within the lumbar spinal cord, excitatory synapses (e.g., proprioceptive afferent synapses involved in the sensory-motor circuitry) that are in contact with PAPs demonstrate a higher level of nanostructure complexity compared to non-contacted synapses (25). In the context of motor neuron neurodegenerative diseases, tripartite synaptopathy has recently been identified as a key hallmark of amyotrophic lateral sclerosis (ALS) disease progression, with ALS murine models (SOD1G93a) and patient human postmortem spinal cord tissue demonstrating a selective loss of PAP contacted synapses (25). With the accumulating evidence of PAPs involvement in synaptic pathologies across other neurological diseases, such as Alzheimer’s disease (26) and Parkinson’s disease (27), this suggests importance of PAPs in healthy synaptic regulation and highlights the vulnerability of tripartite synapses in neurodegenerative diseases. Better understanding of the molecular disease mechanisms mediating intrinsic defects within PAPs could lead to novel therapeutic targeting and treatment development for neurodegenerative diseases.

Proprioceptive afferent synaptic dysfunction is one of the earliest and most consistent phenotype presentations within the disease progression of spinal muscular atrophy (SMA) (28–32), which features the gradual degeneration of motor neuron within the lumbar spinal cord caused by deletion of *SMN1* and global reduction in SMN protein levels (33, 34). Preserving the health and function of these synaptic connections is therefore a critical therapeutic goal for SMA patients. We and others have demonstrated the non-cell autonomous disease mechanisms instigated by astrocytes that exacerbate motor neuron dysfunction in human SMA disease progression, many of which directly relate to the function of PAPs, such as decreased levels of glutamate transporter protein, EAAT1 (35, 36), and abnormal calcium dynamics and response to adenosine triphosphate stimulation (37). Intriguingly, the function of patient-derived motor neurons is improved within direct contact co-cultures featuring healthy-derived astrocytes (35), which suggests that a glial-targeted therapeutic strategy that improves PAP contact-mediated and secreted synaptic support could facilitate motor neuron function in SMA disease pathology. However, targeted SMN restoration within SMA astrocytes only partially restores the disease phenotype (e.g., EAAT1 expression) and has limited improvement on motor neuron function *in vitro* (35). Data from human patients receiving SMN restoration therapies also suggest some ongoing glial pathology and variability in phenotype improvements (38, 39), suggesting there may be additional underlying disease mechanisms that are insensitive to SMN gene augmentation at the point of therapeutic intervention. Hence, developing combined therapeutic approaches targeting SMN expression levels and SMN-independent disease mechanisms may provide greater therapeutic efficacy. While SMN can directly bind to actin and influence the phosphorylation and expression levels of actin binding proteins to influences their activities and cross talk between actin polymerization pathways (40–45), there is evidence to suggest that actin-associated proteins (e.g., PLS3 (46)) and modulating actin signaling pathways (e.g., via ROCK inhibition (47, 48)) can act in an SMN-independent manner to decrease disease severity. Therefore, actin dynamics are likely to be influenced by SMN deficiency but may also provide SMN-independent/related disease mechanisms to target to improve therapeutic outcomes.

The purpose of this study was to further define PAP and motor neuron synaptic defects in human SMA disease pathology towards developing a combination therapy paradigm to improve motor neuron function in SMA. Using microscale cell surface capture mass spectrometry, we characterized the differentially abundant *N-*glycoproteins across healthy and SMA patient-derived astrocytes, which revealed a significant decrease in actin and filopodia-associated proteins within the disease state. In follow-up experiments, we confirmed intrinsic CDC42 actin-based defects within SMA astrocyte filopodia associated with impaired CD44 and phosphorylated ERM (pERM) protein levels. Within our motor neuron astrocyte co-cultures, lower levels of PAP-associated pERM detection were concurrent with a reduced number of synapses in SMA patient-derived cultures compared to healthy samples. Molecular probing of the SMA motor neuron-astrocyte synaptogliosome subcellular fraction also revealed a specific and significant reduction in the levels of the pre-synaptic protein SYN1. Functional recordings of the SMA co-cultures demonstrated motor neuron hyperexcitability at the single cell level (patch clamp) and concurrent diminished synaptic bursting events at the neuronal network level (multi-electrode arrays). In our therapeutic targeting approach, we utilized SMN gene therapy in combination with forskolin, an activator of adenylyl cyclase to increase intracellular cyclic adenosine monophosphate levels, which has been previously used to modulate Rho-family small GTPases activity and actin dynamics. We find that the dual therapy approach provides the most robust improvement in astrocyte filopodia density, synapse formation and motor neuron functional phenotypes, representing a novel treatment paradigm for preserving motor neuron function in SMA disease pathology.

## Methods

### Human stem cell culture and differentiations

The following human induced pluripotent stem cell lines (iPSCs) are detailed in Table 1. iPSC lines were cultured on Corning® Matrigel® Growth Factor Reduced Basement Membrane Matrix (Gibco) coated 6-well plates in Essential 8 medium (Gibco) and were passaged every 3-4 days using Versene (Gibco) when 80% confluent. iPSCs were differentiated into spinal cord-like motor neurons or astrocytes using previously described protocols (35, 49). For co-culture analysis, motor neuron differentiation cultures (Day 14) and astrocyte cultures (P4+) were simultaneously dissociated and combined together in 1:1 motor neuron maturation medium and ScienCell Astrocyte Medium (with 2% B27) at a cell density ratio of 5:1 (coverslips:1.2 x10^5^ motor neurons, 3 x10^4^ astrocytes; 100mm cell culture dishes: 7.2x10^6^ motor neurons 1.8x10^6^ astrocytes; MEA: 2.4 x10^4^ motor neurons, 6 x10^3^ astrocytes).

**Table 1.** Details of human stem cell lines used in this study.

| Line | Sex | Race | Age | SMN1 genotype | Source | Additional information |
| --- | --- | --- | --- | --- | --- | --- |
| Healthy line 1 | Female | Caucasian | 21yrs | Normal | reprogrammed from GM02183 fibroblasts (Coriell); generated in (55) | N/A |
| Healthy line | Female | Caucasian | Not known | heterozygous for deletion of exons 7 and 8 in the SMN1 gene | reprogrammed from GM03814 fibroblasts (Coriell); generated in (55) | Parent of SMA line 2 |
| Healthy line 3 | Male | South Asian | 36yrs | Normal | Greenstone line (GSB-2009) | N/A |
| SMA patient line 1 | Male | Caucasian | 2yrs | Type 1 (SMN2 copy number: 3) | reprogrammed from GM09677 fibroblasts (Coriell); generated in (55) | N/A |
| SMA patient line 2 | Male | Caucasian | 3yrs | Type 2 (SMN2 copy number: 3) | reprogrammed from GM03813 fibroblasts (Coriell); generated in (55) | N/A |
| SMA patient line 3 | Male | Not known | Not known | Type 1 | iPierian | N/A |
| SMA patient line 4 | Female | Asian | 1.95yrs | Type 1 (SMN2 copy number: 2) | Greenstone line (GSB-2945) | N/A |
| SMA patient line 5 | Female | Caucasian | 8yrs | Type 1 (SMN2 copy number: 2) | Greenstone line (GSB-2394) | N/A |

### SMN lentivirus preparation and infection

Lentiviral DNA construct containing FLAG tagged human *SMN1* (hEF1α:SMN-FLAG) was kindly provided by the laboratory of Dr. Xue Jun Li (University of Illinois-Chicago). Lentivirus generation was performed by the Viral Vector Core at the Medical College of Wisconsin/Blood Research Institute; the viral titer for the SMN:FLAG lentivirus used in this study was 7.36E+08 TU/ml. SMA patient-derived astrocytes were transduced at a MOI of 7 with 10ug/ml polybrene for 24 hours. Lentivirus generation of the pLV[Exp]-EGFP-EF1A>hSMN1 vector was outsourced to VectorBuilder. The viral titer for the SMN:GFP lentivirus used in this study was 6.0E+08 TU/ml. SMA patient-derived astrocytes were transduced at a MOI of 3 with 10ug/ml polybrene for 24 hours.

### Microscale cell surface capture mass spectrometry

Astrocytes were plated on to Matrigel coated 6-well plates at a cell density of 1x10^6^ per well. Fresh ScienCell astrocyte medium was applied on to the cells 2hrs prior to beginning protocol. Cells (1 well = 1x sample) were subject to the CellSurfer platform previously developed by the Gundry Lab (50) with some modifications. The oxidation, quenching and biotinylation steps were performed at the Medical College of Wisconsin, after which cells were pelleted, snap frozen and shipped to the University of Nebraska Medical Center for downstream sample processing (lysis, in-solution digestion, peptide quantification, 250 µg of total peptide used for glycopeptide enrichment via streptavidin beads, peptide deglycosylation via PNGaseF, peptide cleanup and elution) and were analyzed using a Dionex UltiMate 3000 RSLCnano system in line with an Orbitrap Exploris 480 MS. Data were searched with Spectronaut, processed with Veneer (51) for cell surface scoring, and MSstats (52) for statistical analysis, similar to our prior studies (50). Only proteins categorized as high or medium confidence surface proteins by Veneer were considered.

### Cell fractionation: plasma membrane enriched fraction and synaptogliosome fraction

Astrocyte monocultures were plated into Matrigel-coated T75 flasks at a density of 1.125x10^6^ and grown to confluency (3-4 days after plate down). Motor neuron-astrocyte co-cultures were plated on to Matrigel-coated 100mm cell culture dishes at a ratio of 5:1 (7.2x10^6^ motor neurons 1.8x10^6^ astrocytes) and grown for a 2-week period to allow synapses to form. To isolate the plasma membrane fraction (astrocyte monocultures) or synaptogliosome fraction (motor neuron astrocyte co-cultures), cells were scraped and collected in 500ul of sucrose buffer (5 mM Tris-HCl pH 7.4, 1 mM EGTA, 1 mM DTT and 320 mM sucrose). Samples were then passed five times through a 26G needle with 1ml plastic syringe and centrifuged at 1000 x g for 10 mins at 4°C. The supernatant was transferred to an ultracentrifugation compatible 1.5ml tube and kept on ice. The pellet was vortexed in the presence of the original volume of sucrose buffer (500ul) and centrifuged again at 1000 x g for 10 min at 4°C. The supernatant was collected and added to the original supernatant (1ml total volume) and the pellet (P1 fraction) was snap frozen on dry ice. The combined supernatants were centrifuging at 23,000 rpm for 20 min at 4°C in an Optima Max-TL Ultracentrifuge (Beckman Coulter). After centrifugation, the supernatant was discarded, and the pellet was resuspended in 1 ml of wash buffer (5 mM Tris-HCl pH 7.4, 1 mM EGTA and 1 mM DTT), and centrifuged at 23,000 rpm for 30 min at 4°C. The supernatant was discarded, and 80-130ul of lysis buffer (100 mM Tris-HCl pH 8.0, 150 mM NaCl, 5 mM EDTA, 1% Triton X-100) was added to the cell pellet before homogenizing via pipetting; this is the plasma membrane enriched fraction/synaptogliosome fraction. During the ultracentrifugation steps, the same lysis buffer was added to the left-over pellet from the initial centrifugation step (P1 fraction). All solutions contained 1x protease inhibitors to maintain protein integrity throughout the procedure. All samples were stored at -80°C prior to downstream western blot analysis.

### Western blot

Samples protein concentration (mg/ml) was determined via Pierce™ BCA Protein Assay (Thermo Scientific) using BSA standards. Samples (17ug of protein per sample) were boiled in 4x loading buffer (with 1% BME) and loaded into 4-15% Mini-PROTEAN® TGX Stain-Free™ Protein Gels (Bio-Rad). The gel was run at 105V for 90 minutes. Protein samples were then transferred to PVDF membranes (Li-COR) at 105V for 60 minutes, which were dried overnight to facilitate protein crosslinking. After methanol rehydration, membranes were incubated in Revert™ 700 Protein Stain (Li-COR) and imaged using an Odyssey Scanner (Li-COR). After removal of the total protein stain using the Revert™ Destaining Solution (Li-COR), membranes were incubated with Odyssey Blocking Buffer (Li-COR) at room temperature for 1 hour. Primary antibodies were diluted in the blocking buffer with 0.2% Tween-20 and incubated overnight at 4°C. Primary antibodies for immunoblotting were used at the following dilution: mouse anti-SMN (1:1000, BD Biosciences), rabbit anti-GFP (1:1000; Cell Signaling), rabbit anti-synapsin I (1:1000; Sigma Millipore), mouse anti-PSD95 (1:1000; Millipore Sigma), rabbit anti-CD147 (1:1000; Invitrogen), mouse anti-Ezrin (1:1000; Sigma Millipore), mouse anti-Na+K+ ATPase α (1:1000; Santa Cruz). Following 3x washes with 1x TBST, membranes were incubated for 45mins at room temperature in secondary antibody solution (blocking buffer with 0.2% Tween-20, 0.02% SDS, 1:8000 secondary antibody). Secondary antibodies used in this study included IRDye 800/680 CW Donkey anti-mouse and IRDye 800/680 CW Donkey anti-rabbit (Li-COR). Blots were imaged using the Odyssey Scanner and protein band intensities were converted into grayscale and quantified using Image Studio (Li-COR) software. Antibody signal was normalized to the total protein REVERT stain according to manufacturer’s instructions.

### Actin remodeling stimulation and forskolin treatment

Astrocytes were plated at a cell density of 5x10^3^ on to Matrigel-coated 1.5 coverslips and allowed to recover for 24hrs. Cells were incubated in phosphate-buffered saline (PBS) supplemented with 1 mM CaCl_2_, 1 mM MgCl_2_ and 20 mM NaN_3_ for 1hr to instigate rapid ATP depletion and reversible actin filament disassembly (53). The buffer was removed and replaced with fresh astrocyte medium for 30 min to allow for ATP recovery and actin cytoskeleton restoration. For the forskolin treatments, cells were incubated for 10µM forskolin (Sigma) for 1hr 30mins or 10µM forskolin was added during the 30min recovery incubation of the actin stimulation assay. Control conditions (UTX) included a full astrocyte medium change and incubation for 1hr 30mins. During the last 30mins of each treatment, CMDFA Green Cell Tracker dye (Thermo Fisher Scientific; final concentration 1uM) was added to cells to allow for full cell morphology visualization.

### Immunocytochemistry

Astrocytes monolayer coverslips and motor neuron-astrocytes coverslips were fixed in 4% PFA for 10-12mins at room temperature, while astrocyte monolayer coverslips for SMN protein detection were fixed in ice cold 1:1 acetone/methanol for 5 minutes at 4°C. Coverslips were washed three times with 1x PBS and stored in PBS at 4°C. Cells were permeabilized in 0.2% Triton X-100 for 10 minutes, before blocking in 10% normal donkey serum, 0.1%BSA for 1 hour. Cells were incubated overnight in primary antibody solution (0.1% Triton X-100, 5% normal donkey serum, 0.1% BSA) at 4°C. The following primary antibodies were used in this study: mouse anti-SMN (1:200, BD Biosciences), rabbit anti-FLAG (1:400, Sigma), rabbit anti-synapsin I (1:800; Sigma Millipore), mouse anti-PSD95 (1:1000; Sigma Millipore), rabbit anti-PERM (1:400; Cell Signaling), mouse EZRIN (1:600; Sigma Millipore), chicken SYN1/2 (1:1000; Synaptic Systems), mouse anti-SYT1 (1:1000, Synaptic Systems), mouse CD44 (1:500; BD Biosciences). Cells were washed 3x with 1x PBS (5 minutes per wash) before incubating with secondary antibodies at room temperature for 1 hour. Secondary antibodies used in this study included donkey anti-mouse Alexa Fluor 488 Plus (Invitrogen), anti-rabbit Alexa Fluor 568 (Invitrogen) and anti-chicken Alexa Fluor 647 Plus (Invitrogen). iPhalloidin-647 (Abcam) was used to visualize F-actin. Primary antibodies were omitted from secondary only controls. Cells were washed 3x with 1x PBS (5 minutes per wash) prior to 4’,6-diamidino-2-phenylindole (DAPI; 1:3000; Thermo Fisher Scientific) incubation for 5 minutes at room temperature to stain nuclei. Cells were washed 3x with 1x PBS (5 minutes per wash) and were mounted onto Superfrost Plus microscope slides (VWR) in VECTASHIELD antifade mounting medium (Vector Laboratories). Coverslips were sealed with nail polish and stored at 4°C prior to imaging.

### Confocal microscopy

Z-projection images were acquired of astrocyte-stained coverslips using the Zeiss LSM980 confocal laser scanning microscope with AiryScan 2 and ZEN 3.4 (blue edition) acquisition software (Zeiss). Images were processed using ZEN 3.4 and ImageJ software.

### Filopodia density quantification

ImageJ plugin tool, FiloQuant (54), was used to quantify astrocyte filopodia density. Z-projection confocal images containing CMDFA Green Cell Tracker dye signal were loaded into the Imaris software. Signal was changed to grey scale and a 0.4 µm surface was applied to mark dye signal to create a masking channel where signal within cells is transferred onto a pure black background. This facilitated the accurate identification of filopodia by the FiloQuant tool. Area of interest was demarcated, and the following thresholds were used in the filopodia analysis: cell edge threshold = 3 and filopodia threshold = 20. The tagged skeleton RGB images containing the original image with quantified filopodia regions highlighted in magenta were used as representative images in main text figures. The total number of filopodia was divided by the sum of the total cell edge length (microns) to give a normalized filopodia quantification value (i.e., filopodia density).

### CDC42-GTP G-LISA assay

The colorimetric CDC42 G-LISA GTPase activation assay kit (Cytoskeleton, Inc) was utilized to assess the amount of active CDC42-GTP within healthy and SMA astrocyte cultures at baseline and after actin stimulation via the ATP depletion and recovery assay. The protocol was followed according to manufacturer’s instructions. Briefly, astrocytes were collected and homogenized in ice cold lysis buffer containing protease inhibitors and centrifuged at 10,000 x g for 1mins at 4°C before collecting cell lysate and storing at -80°C. Samples protein concentration (mg/ml) was determined via Pierce™ BCA Protein Assay (Thermo Fisher Scientific) using BSA standards, and prior to running assay, samples were diluted to 1mg per sample. Samples were added to CDC42-GTP G-LISA plate alongside CDC42 control protein samples and agitated at 4°C for 15mins, before proceeding to the anti-CDC42 primary antibody (1:80 dilution), secondary HRP antibody (1:100), HRP detection and HRP stop buffer steps prior to measuring absorbance at 490nm. In between step, the plate was washed thoroughly with wash buffer and antigen presenting buffer was added prior to the primary antibody step.

### Imaris software analysis

The full suite of Imaris modules was made available through the MCW OxCAM-EM core. This study made use of the *Imaris Core* (render 3D/4D images, detect objects, capture snapshots and animations), *Imaris MeasurementPro* (report and interact with detected object measurements) and *Imaris Coloc* (visualize and quantify colocalized regions), and *Imaris Batch* (utilize saved protocols for batch analysis tools). A 0.4 µm surface was applied on to green dye signal to generate surface for astrocyte morphology visualization. The Spots tool was used to label SYN1 and PSD95 puncta (1µm, ≤0.5µm distance between spots was determined as a synapse) and pERM puncta (0.6µm) for synapse and predicted PAP quantification, respectively. The Spots tool was also used to label and quantify pERM and CD44 puncta (1µm, ≤0.5µm distance between spots was determined as colocalized signal) in astrocyte filopodia regions. Surface and spot tool parameters were generated and applied to all images for each analysis using the batch tool.

### Structured illumination microscopy

The first-generation Nikon N-SIM composed of a fully motorized Nikon Ti, an *Andor iXon EMCCD camera and* 4 lasers (405/488/540/633 nm) was used in this study. The 100x SR oil objective was used to acquire images of CD44, pERM, EAAT1 and Phalloidin signal in astrocyte filopodia. The microscope is run by the Nikon’s Advanced Research package of Nikon Elements and images were acquired in multi-dimensional acquisition mode (3D SIM, 0.2um step multi-position, z-projection stack approx. 4.4 µm). Images were saved as ndi. files and were imported into the Imaris software for reconstruction.

### Voltage clamp electrophysiology

Coverslips were transferred to a recording chamber perfused with artificial cerebrospinal fluid (ACSF) containing (in mM): 127 NaCl, 1.9 KCl, 1.2 KH_2_PO_4_, 2.2 CaCl_2_, 1.4 MgSO_4_, 26 NaHCO_3_, and 10 glucose. The solution was continuously bubbled with 95% O_2_ and 5% CO_2_. Whole-cell patch-clamp recordings were performed using a Multiclamp 700B amplifier under infrared differential interference contrast (IR-DIC) microscopy. Data acquisition and analysis were conducted with a DigiData 1440A digitizer and pClamp 11 software (Molecular Devices). Signals were filtered at 2 kHz and sampled at 10 kHz. Glass recording pipettes (3–5 MΩ) were filled with an internal solution containing (in mM): 140 K-gluconate, 5 KCl, 2 MgCl_2_, 10 HEPES, 0.2 EGTA, 4 Mg-ATP, 0.3 Na_2_GTP, and 10 Na_2_-phosphocreatine at pH 7.2 (with KOH). Motor neurons were initially recorded in voltage-clamp mode. Cell capacitance was determined by applying brief voltage steps and analyzing the resulting capacitive transients. Recordings were then switched to current-clamp mode. Step current injections (-40 to 100 pA, 10 pA increments, 500 ms) were applied to measure action potentials and input resistance (R_in_). Rheobase was determined by applying depolarizing current steps in 5 pA increments until the first action potential was evoked. R_in_ was calculated from the slope of the linear portion of the current–voltage (I-V) relationship. Resting membrane potential (RMP) was recorded concurrently at I = 0 pA. All recordings were performed at 32 ± 1°C using an automatic temperature controller (Warner Instruments, Hamden, CT). Series resistance (15-30 MΩ) was monitored throughout, and data were excluded if resistance changed by >20%.

### Multi-electrode array

Motor neurons and astrocytes were plated on to poly-L-lysine (Sigma)/laminin (Sigma) coated 48 well CytoView microelectrode array (MEA) plates (Axion Biosystems) in a 5µL droplet directly over the recording electrodes of each well. Cells were left to attached for 1 hour at 37°C before carefully flooding the wells with 300µl of media. Base medium or medium containing 10uM forskolin was replaced 3 times per week. Spontaneous and electrically evoked motor neuron activity was recorded every other day for approximately 1 month starting from Day 7 post-plate down. MEA recordings were performed by the 1^st^ generation Maestro system (Axion Biosystems). Cells were rested for 10 minutes on the pre-heated (37°C) MEA stage prior to starting the 6-minute recordings. Version 2.4.2.13 of the AxIS acquisition software (Axion Biosystems) was used to record activity across 16 electrodes per well with a sampling frequency of 12.5 kHz and a digital high pass filter of 5Hz IIR. The spike-detecting threshold was defined as voltage exceeding 6 standard deviations away from the mean background noise. Butterworth digital filter settings with high (200 Hz) and low (3kHz) pass cut off frequencies were applied during recordings. 5 spikes/minute was the threshold used to determine an active electrode. For electrically evoked recordings, the Neural Stimulation with Artifact Elimination (0.5V for 400µs (36400µs total operation time) was applied to the cells by stimulating 1 electrode per well once every 10 seconds. Statistic Compiler files and Spike Detector files were extracted from raw AxIS (spontaneous recordings) and raw AxIS Artifact Eliminator files (electrically evoked recordings) for downstream analysis of weighted mean firing rate and bursting activity. The NeuralMetricTool (Axion Biosystems) was used to generate raster plots.

### Statistics and rigor

The number of biological lines (N) and experiment replicates (n) are stated in the legend for each figure. Statistical significance within the proteomics data was determined as adjusted p-value (-log_10_P) ≤0.05; log2 fold change ≤ or ≥1. All statistical analysis and generation of graphs was performed in GraphPad Prism software. One-way ANOVA with Bonferroni multiple comparison testing, Two-way ANOVA with Bonferroni multiple comparison testing or unpaired t-test statistical testing was performed in this study. Statistical significance was determined as p-value<0.05.

## Results

While SMN is preferentially localized to nuclear gems and the cytoplasm, SMN can also localize to plasma membrane protrusions during states of active actin remodeling (56). We performed a subcellular fractionation protocol to enrich for the plasma membrane fraction within our stem cell-derived astrocyte samples. Western blot analysis confirmed the enrichment of established glial cell surface proteins (CD147, Na+K+ ATPase α) within the plasma membrane fraction compared to the whole cell lysate samples (Fig. 1A). We also confirmed the presence of SMN protein within the plasma membrane fraction, which was depleted in SMA patient-astrocyte samples compared to healthy control samples (Fig. 1A). To assess how SMN deficiency influences the astrocyte cell surfaceome, we utilized microscale cell surface capture mass spectrometry to quantitatively compare plasma membrane localized *N-*glycoproteins in healthy and SMA patient-derived astrocytes. Principal component analysis reveals a significant separation and clustering of samples (PC1: 37.5%, based on disease state (i.e., healthy vs SMA; Fig. 1B)). We identified a total of 276 cell surface *N-*glycoproteins, of which one was uniquely observed in SMA (*e.g.,* NAALAD2) and two were uniquely observed in control (e.g., FGFR3, L1CAM). Of the 273 observed in both conditions, 81 (29%) were significantly different in abundance between conditions (adjusted p-value (-log_10_P) ≤0.05; log2 fold change -1≤ or ≥1); 44 proteins demonstrated higher abundance in patient-derived samples, while 37 proteins showed significantly reduced levels in SMA patient-derived astrocytes compared to healthy samples (Fig. 1C). Notably, several proteins with the most significant increase in abundance in SMA astrocyte samples compared to control samples are involved in inflammatory responses (COLEC12, PTPRO, IL17RD), and astrocyte reactivity (ICOSLG) (Fig. 1D), while cell surface proteins most significantly decreased are associated with regulating actin cytoskeleton dynamics required for filopodia motility (TENM2, CD44, CEMIP2), infiltration of astrocyte processes into the neuropil (TENM4), and synaptic plasticity (TNC) (Fig. 1E). Together, the cell surfaceome data suggests that SMN deficiency contributes to astrocyte reactivity, which is a consistent previously observed phenotype (37, 57, 58), but may lead to actin remodeling defects, particularly those involved filopodia outgrowth and retraction.

**Figure 1.**
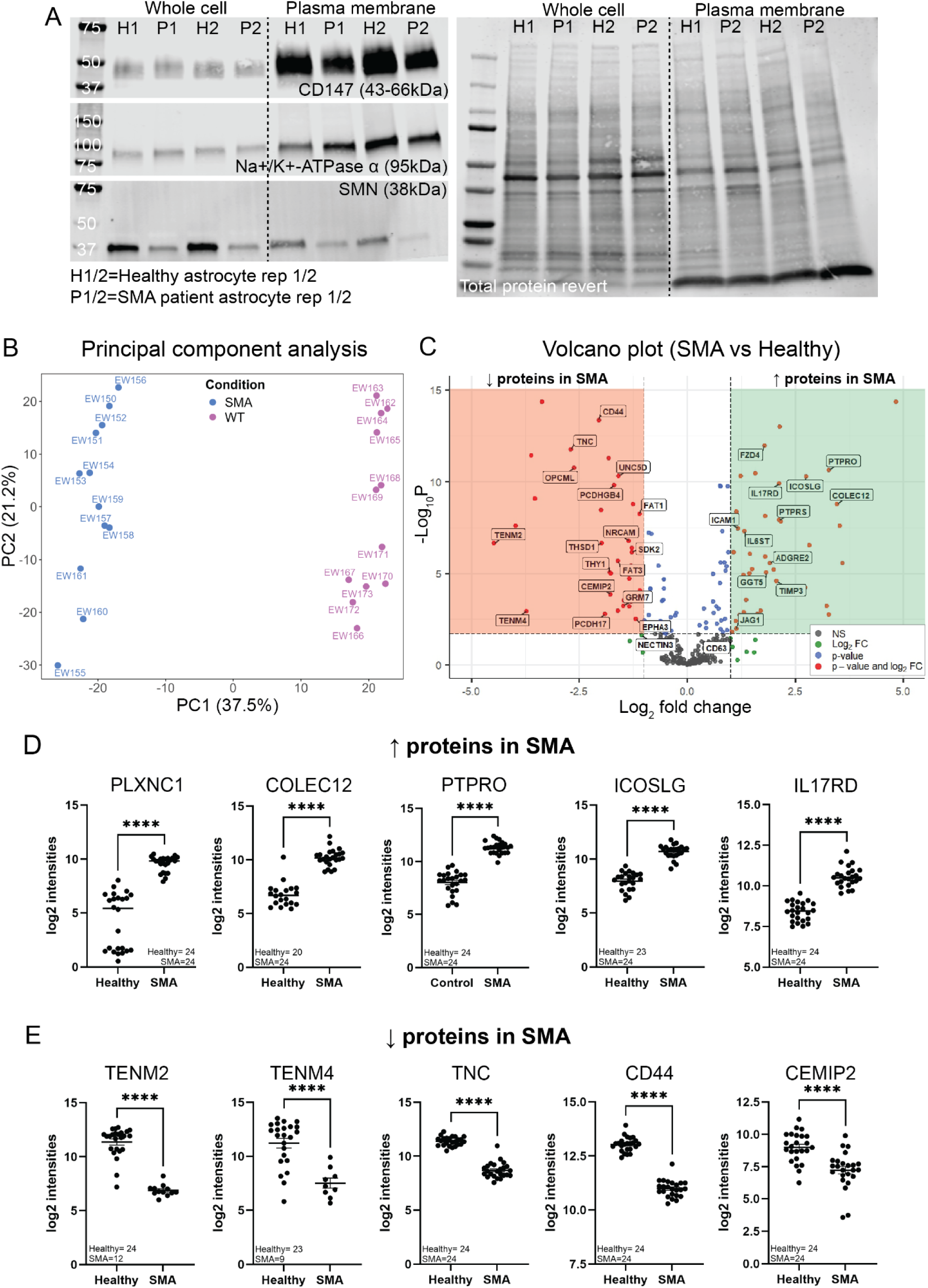
Differentially abundant cell surface proteins in healthy vs SMA patient-derived astrocytes. **(A)** Western blot analysis of astrocyte cell surface protein levels (CD147 and Na+K+ATPase α) and SMN protein levels in whole cell lysates and plasma membrane enriched lysates derived from healthy and SMA patient-derived astrocyte samples. Microscale cell surface capture mass spectrometry was performed on healthy and SMA patient-derived astrocyte samples. **(B-E)** Summary of cell surface capture mass spectrometry data. **(B)** Principal component analysis assessing sample variability and clustering by condition. **(C)** Volcano plot demonstrates quantified cell surface N-glycoproteins that are differentially abundant (-log10 adjusted p-value <0.05; log2FC>1) in healthy and SMA patient astrocytes. Individual log2 intensity plots are shown for proteins that had lower **(D)** or higher **(E)** abundance in SMA patient-derived astrocytes. Unpaired t-tests; ****p≤0.0001. N=2 (biological replicates), n= 12 (technical replicates)

We next assessed the filopodia dynamics of healthy and SMA patient-derived astrocytes at baseline and after actin stimulation via an ATP depletion (via 20mM sodium azide treatment) and recovery assay (NaN3 + R). Fig. 2A shows representative confocal images of healthy and SMA astrocytes labelled with a CMFDA-based dye that was used to quantify filopodia density (number of filopodia/cell perimeter) using the FiloQuant ImageJ plugin tool (54). At baseline (UTX), SMA patient-derived astrocytes demonstrated a significant decrease in filopodia density compared to healthy astrocytes (Fig. 2B). Acute actin remodeling stimulation (NaN3 +R) a significant increased filopodia density specifically within healthy astrocyte samples compared to baseline, while SMA astrocyte filopodia density remained equivalent between baseline and NaN3 + R conditions (Fig. 2B). The Imaris surface tool was used to visualize cell morphology and filopodia of individual cells, which highlights the reduced number of filopodia of SMA patient-derived astrocytes compared to healthy astrocytes at baseline and after actin remodeling stimulation (NaN3 + R) (Fig. 2C). Quantification of the active form of CDC42 (CDC42-GTP) responsible for initiating actin remodeling within filopodia demonstrated no significant differences between healthy and SMA astrocytes at baseline (Fig. 2D; p=0.5129). However, post actin remodeling stimulation (NaN3 + R), CDC42-GTP levels showed an increasing trend within healthy astrocytes compared to baseline, while levels of CDC42-GTP remained unchanged within SMA patient-derived astrocytes (Fig. 2D). Together, this suggests an intrinsic defect in CDC42-associated actin dynamics within filopodia of SMA astrocytes.

**Figure 2.**
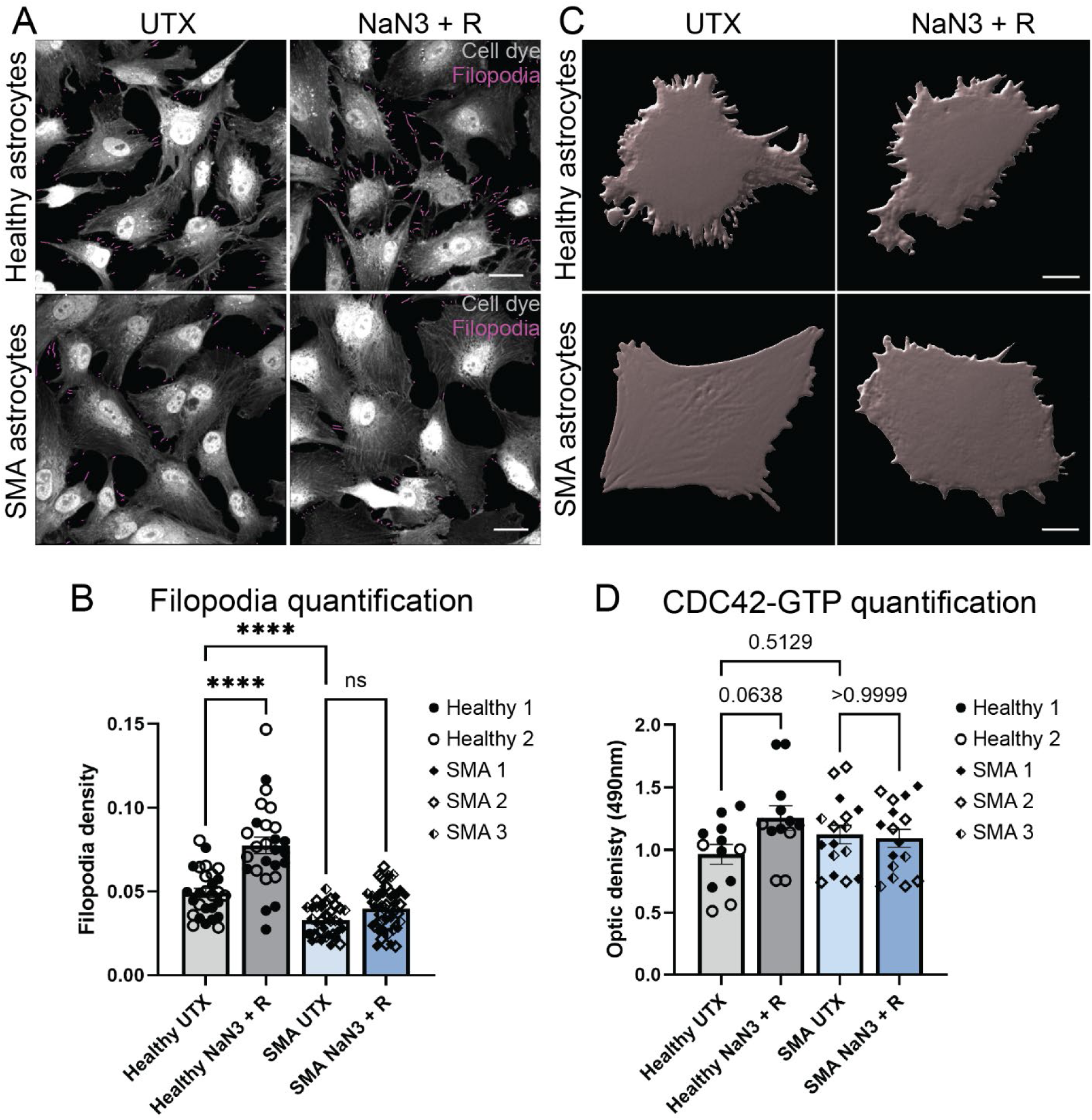
SMA patient astrocytes demonstrate decreased filopodia density and CDC42-GTP levels. **(A)** Representative images from FiloQuant analysis used to quantify filopodia density in healthy and SMA patient astrocyte monocultures at baseline and after stimulating actin remodeling using the ATP depletion and recovery assay (NaN3 +R). CMFDA dye was used to visualize the cells (grey signal) and filopodia quantified by FiloQuant tool are highlighted in magenta. Scale bar: 30µm. **(B)** Imaris surface and mask tool applied on to individual healthy and SMA patient astrocyte cells at baseline and after ATP depletion and recovery assay. Scale bar: 15µm. **(C)** Quantification of filopodia density from FiloQuant analysis across baseline and treatment conditions for healthy and SMA patient astrocyte samples. One-way ANOVA with Bonferroni multiple comparison statistical testing; ****p<0.0001. **(D)** Optic density readout from CDC42-GTP G-LISA assay measuring the activated form of CDC42 across baseline and treatment conditions for healthy and SMA patient astrocyte samples. One-way ANOVA with Bonferroni multiple comparison statistical testing, p-values not statistically significant. N=5 (biological replicates), n= 4 (technical replicates).

We further investigated the molecular mechanism contributing to filopodia actin remodeling defects within SMA patient-derived astrocytes. Our mass spectrometry data reveal significantly lower relative abundance of hyaluronic acid (HA) receptor (CD44) and regulator of HA metabolism (CEMIP2) in SMA astrocytes compared to healthy astrocytes, suggesting SMA astrocytes may have abnormal CD44-hyaluronic acid signaling, which plays an important role in regulating the actin cytoskeleton via RhoA GTPase pathways (i.e., CDC42) to influence filopodia dynamics. This interaction is facilitated by phosphorylated ERM proteins (pERM), which help to stabilize the CD44-hyaluronic acid interactions and act as linker proteins bringing CD44 and actin filaments in close proximity at the plasma membrane (17–19). Immunocytochemistry analysis and confocal microscopy confirmed lower levels of CD44 protein and revealed a significant decrease in pERM protein signal within SMA astrocytes compared to healthy astrocytes (Fig. 3A&B). Notably, both CD44 and pERM are highly enriched within the filopodia of healthy astrocytes (Fig. 3A). Additionally, healthy astrocytes also typically demonstrated a smaller cell size with thinner processes and prominent filopodia, while SMA patient-derived astrocytes showed a larger, swollen appearance with distinctive actin stress fibers as shown through phalloidin staining (Fig. 3A&B). Structured illumination microscopy of astrocyte filopodia allowed for better visualization of the CD44 and pERM proteins, which confirmed the lower levels of CD44 and pERM protein signal in SMA astrocytes compared to healthy samples at baseline and after stimulating actin remodeling (Fig. 3C). Of note, the PAP-associated protein and glutamate transporter EAAT1 was readily detected within these filopodia regions within healthy astrocytes, while low levels were detected in SMA astrocyte filopodia (Fig. 3D). Using the Imaris spot tool function, we quantified the CD44 and pERM puncta within astrocyte filopodia (Fig. 3E) which revealed a statistically significant decrease in pERM (Fig. 3F), CD44 (Fig. 3G), and co-localized CD44/pERM (Fig. 3H) expression within SMA patient-derived astrocytes cultures compared to healthy astrocytes at baseline and after actin remodeling stimulation.

**Figure 3.**
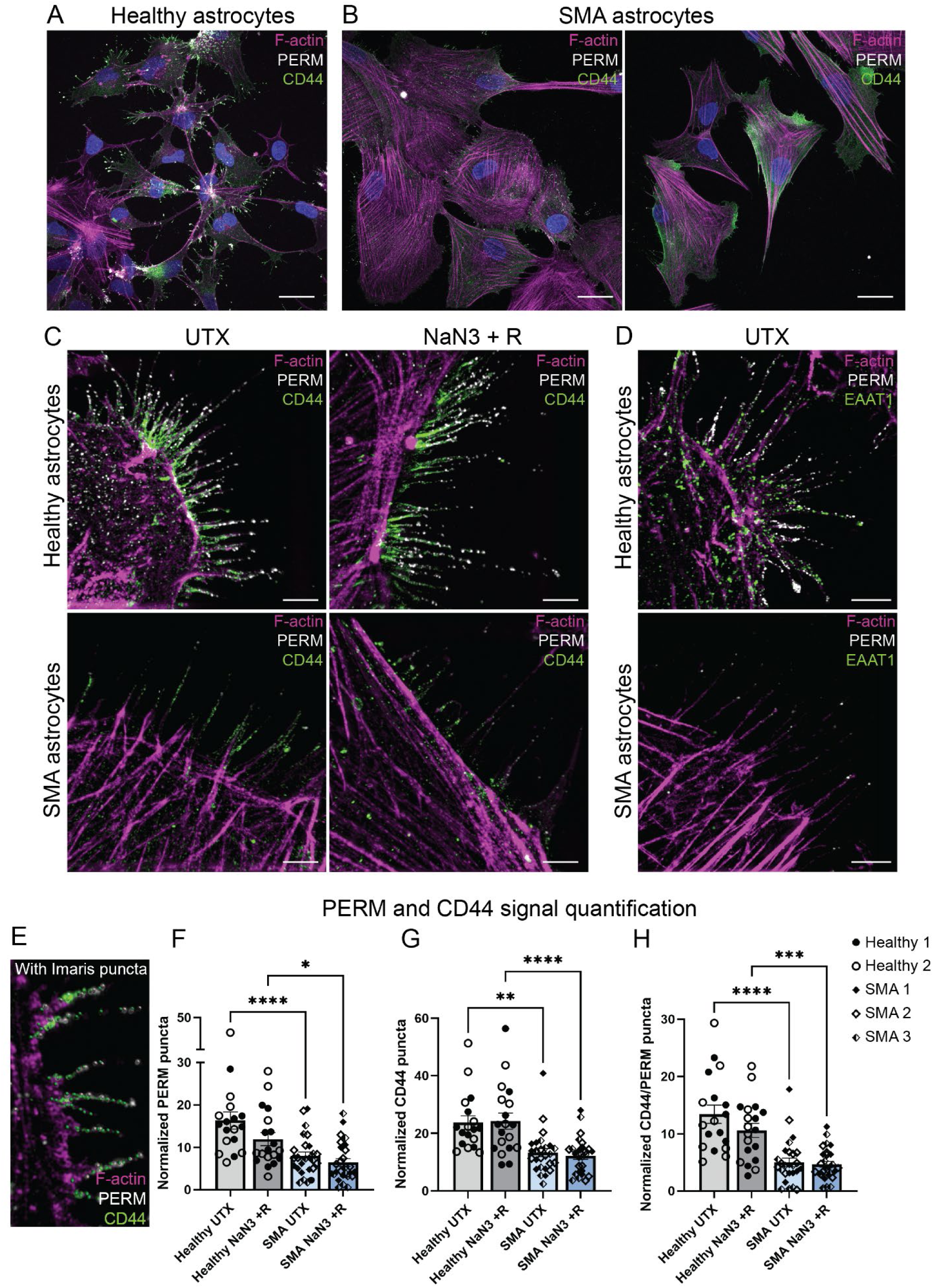
SMA patient astrocyte filopodia demonstrate decreased levels of PAP-associated proteins, CD44 and phosphorylated ERM. Representative confocal images demonstrating CD44 (green signal) and PERM (grey signal) expression levels and localization within **(A)** healthy and **(B)** SMA patient astrocytes. Scale bars: 30µm. **(C)** Representative structured illumination microscopy images of CD44 (green) and PERM (grey) signal within healthy and SMA patient astrocyte filopodia at baseline and after ATP depletion and recovery assay (NaN3 +R). Scale bars: 4µm. **(D)** Representative structured illumination microscopy images of glutamate transporter, EAAT1, (green) and PERM (grey) signal within healthy and SMA patient astrocyte filopodia at baseline. Scale bars: 10µm. **(E)** Example of Imaris software puncta identification CD44 and PERM fluorescent signal within astrocyte filopodia. Puncta quantification of **(F)** PERM, **(G)** CD44, **(H)** co-localized CD44/PERM (distance between puncta ≤0.5µm) signal in healthy and SMA patient astrocyte filopodia at baseline and after ATP depletion and recovery assay (NaN3 +R). Puncta number was normalized to the number of filopodia observed per image. One-way ANOVA with Bonferroni multiple comparison statistical testing; ****p<0.0001, ***p=0.0002, **p=0.0011, *p=0.0131. F-actin was visualized throughout images using AF647-conjugated phalloidin (magenta signal). N=5 (biological replicates), n= 3 (technical replicates).

Our next step was to determine a therapeutic strategy to improve the filopodia-associated actin phenotype observed within SMA patient-derived astrocytes. Previous work has shown that increasing SMN expression in astrocytes provides a partial improvement in SMA astrocyte phenotypes (35, 57). Therefore, we combined the SMN1 gene therapy approach with forskolin, an adenylyl cyclase activator that leads to an increases cyclic AMP and activation of a filopodia-associated actin remodeling cascade via the CDC42 pathway (59, 60) (Fig. 4A). We initially tested this treatment paradigm within the astrocyte monocultures to assess how SMN augmentation via lentiviral expression of SMN and/or forskolin treatment influences filopodia dynamics of SMA patient-derived astrocytes. Previous work has demonstrated that forskolin exposure can increase SMN protein accumulation and stability with prolonged treatment (24hrs+) (61). However, we show that acute forskolin treatment (10µM; 1hr 30min treatment) does not increase SMN protein levels in SMA and SMA SMN re-expressing astrocyte monocultures (Fig. 4B&C), suggesting cell responses are likely to occur independently of SMN protein levels. Healthy astrocytes treated with forskolin at baseline (F) or during the recovery period after actin remodeling (NaN3 + R(F)) demonstrated increased filopodia density (Fig. 4D&E), equivalent to the higher levels observed in healthy astrocyte cultures after actin stimulation (NaN3 + R) (Fig. 4E). Intriguingly, this same trend and similar levels of increased filopodia density were demonstrated by SMA patient-derived astrocytes with forskolin treatment at baseline and during the recovery period after actin remodeling (Fig. 4D&E). Of note, while the filopodia density was increased after forskolin application, the overall SMA astrocyte cell morphology still appeared larger and swollen compared to healthy astrocytes (Fig. 4D). At baseline (UTX), SMA astrocytes transduced with either lenti SMN:FLAG or lenti SMN:GFP demonstrated increased filopodia density compared to untreated SMA astrocytes and equivalent filopodia density levels compared to healthy astrocytes at baseline (Fig. 4F-I). Post-actin stimulation (NaN3 + R) and after forskolin treatment (F), SMN:FLAG SMA astrocytes maintained filopodia density compared to SMN:FLAG SMA astrocyte untreated conditions (Fig. 4F&G); however these levels were still higher compared to baseline (UTX) and actin stimulated (NaN3 +R) SMA astrocyte filopodia density (Fig. 4G). SMN:GFP SMA astrocytes demonstrated a significant increase in filopodia density after actin stimulation (NaN3 + R) and an increased trend in filopodia density after forskolin treatment (Fig. 4H&I), which were also at higher levels compared to SMA astrocyte conditions and at equivalent levels of filopodia density achieved by healthy astrocyte after actin stimulation (Fig. 4I). Across the forskolin treated cultures, it was also noted that SMN re-expressing SMA astrocytes often demonstrated the formation of thicker cellular branches and an elongated cell shape, similar to the appearance of healthy astrocytes treated with forskolin (Fig 4.D, F &H). The addition of forskolin during the recovery period of the actin stimulation assay (NaN3 +R(F)) resulted in striking morphology changes in SMN re-expressing SMA astrocytes, which demonstrated significant branching and stellation compared to cells at baseline (Fig. 4F&H). Quantification analysis revealed a significant increase in filopodia density across SMN:FLAG and SMN:GFP SMA astrocytes compared to baseline (UTX) cells, reaching equivalent filopodia density levels to those observed in healthy astrocyte NaN3 +R(F) conditions (Fig. 4G&I). Overall, these data suggest that forskolin and SMN protein levels can independently contribute to improving filopodia dynamics in SMA astrocytes, but co-administration can dramatically improve astrocyte filopodia and cell morphology during periods of active actin remodeling.

**Figure 4.**
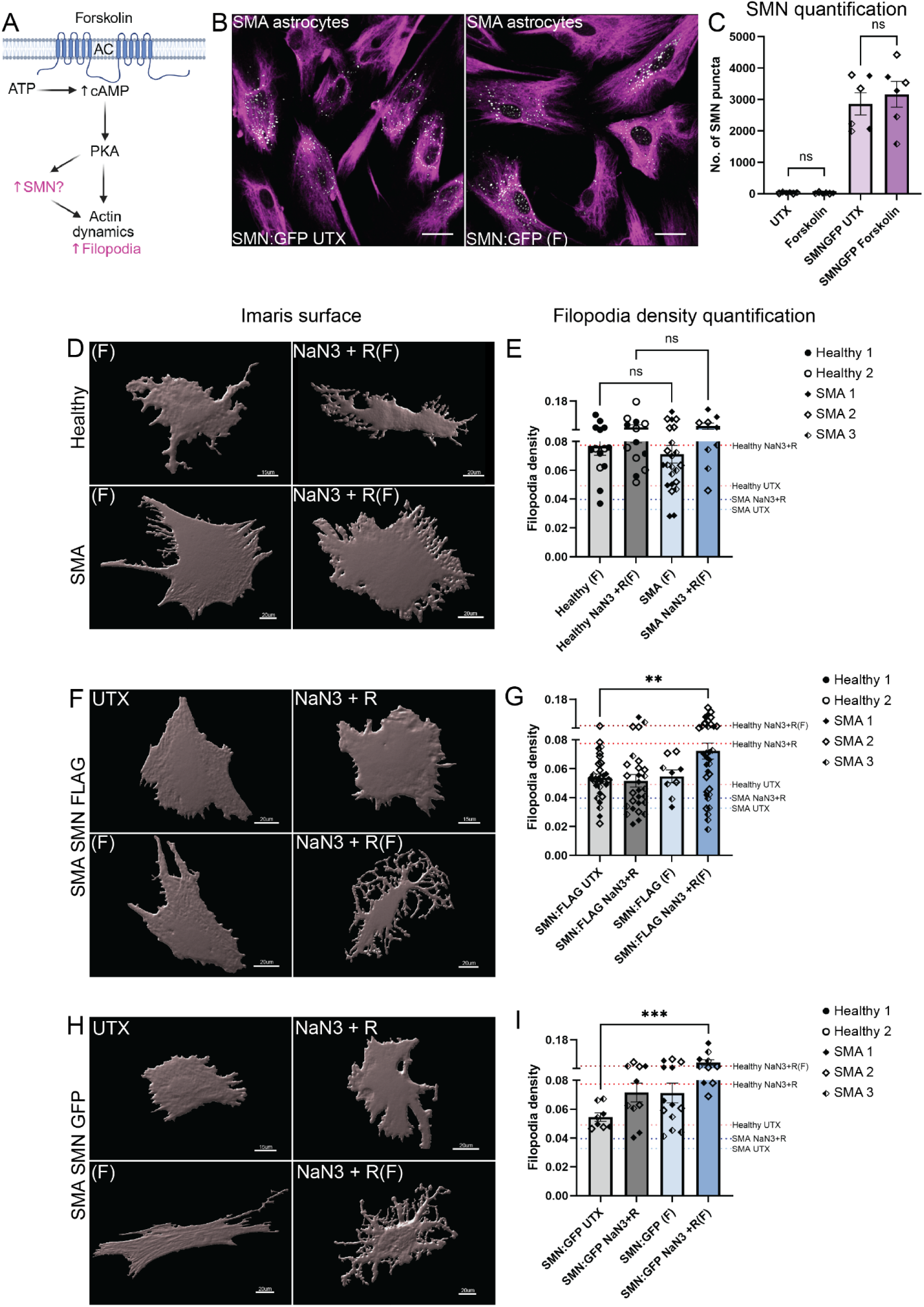
Forskolin improves SMA astrocyte filopodia density at baseline, with SMN re-expression and during actin remodeling. **(A)** Schematic depicting downstream cellular mechanism activated by forskolin treatment. Forskolin application activates adenylyl cyclase (AC) at the plasma membrane, leading to an increase in intracellular levels of cyclic adenosine mono-phosphate (cAMP) and downstream activation of the protein kinase A (PKA) pathway. This can modulate SMN protein levels via phosphorylation and stabilization, in addition to actin remodeling via activation of CDC42. **(B)** Representative confocal images of SMN signal (grey) in SMN re-expressing SMA astrocytes at baseline (SMN:GFP UTX) and after 10µM forskolin treatment (1hr 30min incubation; SMN:GFP (F). VIMENTIN signal (magenta) was used to visualize astrocytes Scale bars: 30µm. **(C)** Quantification of SMN puncta in SMA patient astrocytes and SMN re-expressing SMA astrocytes and after 10µM forskolin treatment. Unpaired t-test; ns=not significant. Imaris surfaces outlining cell morphology **(D)** and quantification of filopodia density **(E)** for healthy and SMA astrocytes after forskolin treatment and forskolin treatment added during recovery period of actin remodeling (NaN3 + R(F). Imaris surfaces outlining cell morphology and quantification of filopodia density for SMN:FLAG re-expressing SMA astrocytes **(F&G)** and SMN:GFP re-expressing SMA astrocytes **(H&I)**. Conditions include baseline (UTX), forskolin treatment (F), actin remodeling via ATP depletion and recovery assay (NaN3 + R) and forskolin treatment added during recovery period of actin remodeling (NaN3 + R(F). Average filopodia density of healthy and SMA astrocyte datasets (featured in Figure 1) is depicted as threshold dotted lines in graphs. One-way ANOVA with Bonferroni multiple comparison statistical testing; **p=0.0082, ***p=0.0005. N=5 (biological replicates), n= 3 (technical replicates)

We next utilized motor neuron-astrocyte co-cultures to assess if these intrinsic defects in actin-associated filopodia dynamics of SMA patient-derived astrocytes influences PAP and motor neuron synapse formation. We isolated PAPs from healthy and SMA patient-derived motor neuron astrocyte co-cultures using a sucrose-based fractionation method (Fig. 5A) and then performed immunoblotting on non-enriched (P1) and synaptogliosome enriched (SG) fractions to probe for pre- (SYN1) and post- (PSD95) synaptic terminal proteins and PAP-associated protein CD147 (62) and EZRIN (8) (Fig. 5B). For these analyses, we included additional samples derived from healthy (Line 3; male) and SMA patient (Line 4 and 5; female; SMA Type I patients) iPSC lines to assess if sex-associated synaptic phenotypes could also be observed. All healthy and SMA SG fractions showed a significant enrichment in SYN1 and PSD95 (Fig. 5C) and CD147 (Fig 5D) compared to P1 fractions. While not statistically significant, we observed a trending enrichment of EZRIN in healthy SG samples versus P1 fractions, but not in the SMA samples (Fig. 5D). We attempted but failed to successfully obtain phosphorylated EZRIN signal via immunoblotting due to antibody detection issues (data not shown). When comparing proteins levels across SG fractions, we noticed a striking decrease in SYN1 protein levels specifically within male SMA patient-derived SG samples compared to healthy SG samples, while SYN1 protein levels remained high in female SMA patient-derived SG fraction (Fig. 5E). PSD95 CD147, and EZRIN remained unchanged across healthy and SMA SG samples (Fig. 5E&F). Interestingly, we detected SMN protein within the SG samples across all samples (Fig. 5G), although it was not significantly enriched within this fraction. We confirmed that all male and female SMA patient-derived samples had lower levels of SMN protein within the SG fraction compared to the equivalent healthy samples (Fig 5F). By molecularly probing the synaptogliosome fraction of SMA patient-derived samples, our data suggest that SYN1, a pre-synaptic terminal protein involved in neurotransmitter release is significantly affected in SMA disease pathology, particularly within the male samples.

**Figure 5.**
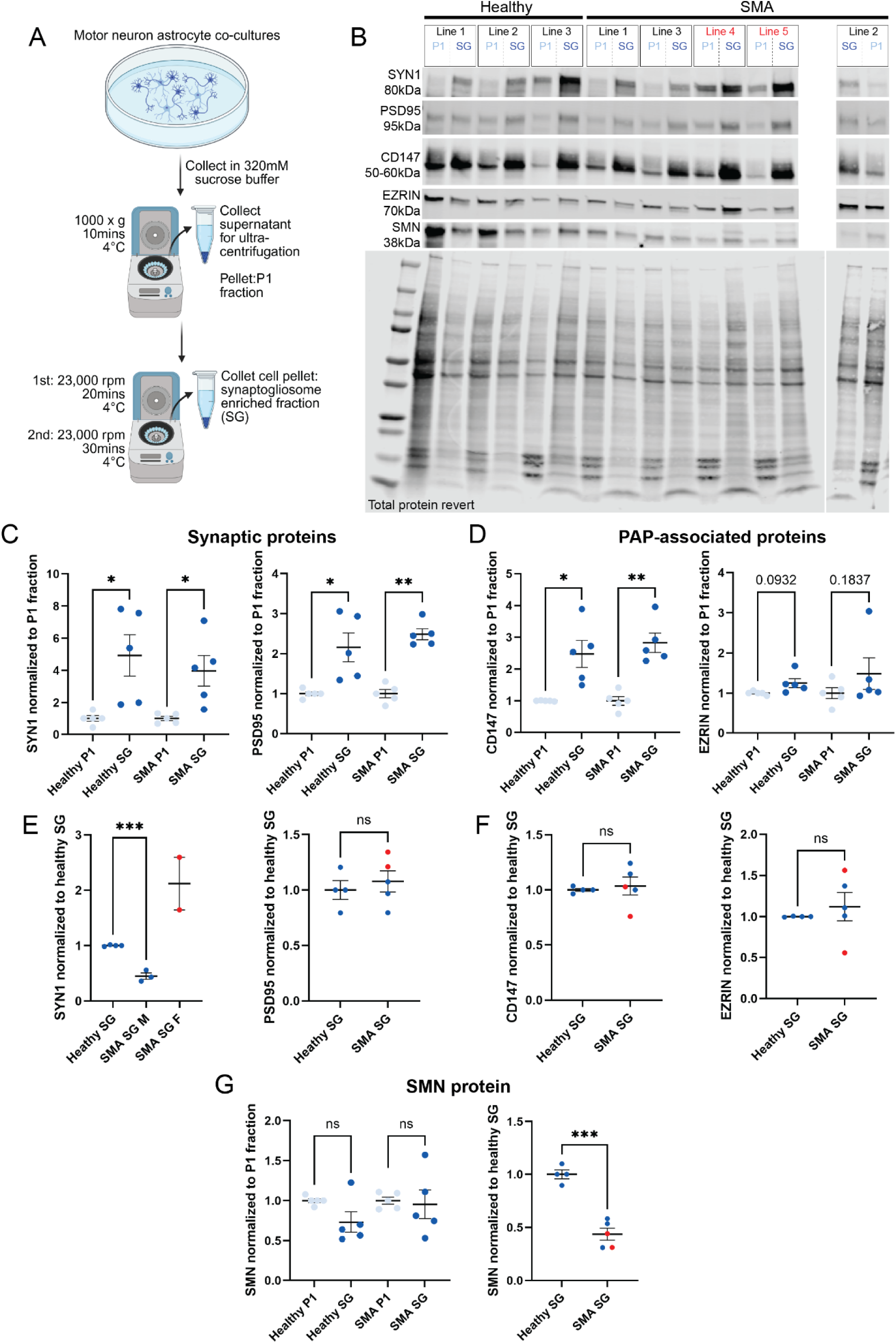
Synaptogliosomes from SMA patient-derived motor neuron astrocyte co-cultures demonstrate decreased levels of synapsin I protein. **(A)** Schematic of synaptogliosome (SG) isolation protocol. **(B)** Immunoblots of P1 and SG fractions from iPSC-derived healthy and SMA patient motor-neuron astrocyte co-cultures demonstrating SMN, synaptic (SYN1, PSD95) and PAP-associated (CD147, EZRIN) protein signal. All protein signal was normalized to total protein revert signal. Quantification (fold change enrichment) of the synaptic **(C)** and PAP-associated **(D)** proteins within the SG fraction compared to the P1 fraction. Quantification (protein level changes) of the synaptic **(E)** and PAP-associated **(E)** proteins between healthy and SMA SG fractions. **(G)** Quantification and protein level changes) of SMN signal within the SG fraction compared to the P1 fraction (fold change enrichment) and between healthy and SMA SG fraction (protein level changes). Paired t-test: P1 vs SG protein signal; Unpaired t-test: healthy SG vs SMA SG protein signal. *p<0.05, **p<0.003.; ***p=0.0001. N=8 (biological replicates), n= 1 (technical replicate).

Subsequently, we applied the astrocyte-targeted SMN1 gene therapy and forskolin treatment paradigm to the SMA patient-derived co-cultures (Fig. 6A-C). We used a prolonged forskolin treatment (2-week period, replenished every 2 days) to overlap with synapse formation and maturation occurring within the culture system. We confirmed a significant increase in SMN levels and detection of GFP within the P1 fraction of SMN:GFP expressing astrocyte co-culture samples (Fig. 6D). Compared to untreated conditions, forskolin treated SMA co-cultures did not show increased SMN levels; however, SMA co-cultures containing SMN re-expressing astrocytes with forskolin treatment demonstrated significantly higher levels of SMN compared to SMA co-cultures containing SMN re-expressing astrocytes without forskolin treatment (Fig. 6D). Similar trends were observed within the SG samples, with forskolin treated SMN:GFP expressing astrocyte co-cultures showing higher levels of SMN compared to healthy baseline samples (Fig. 6D); however, these were not statistically significant. In forskolin treated SMA SG samples, we noted a significant improvement in SYN1 protein levels that were equivalent to healthy control levels, particularly for the male SMA patient-derived samples, compared to SYN1 levels within SMN:GFP expressing astrocyte SMA co-cultures samples without forskolin treatment (Fig. 6E). Levels of PSD95 across all SMA SG samples remained similar to healthy SG levels (Fig. 6F). EZRIN levels appeared equivalent to healthy samples in SMN:GFP expressing SMA astrocyte co-culture samples, while forskolin treated samples demonstrated a minor reduction in EZRIN levels (Fig. 6G). Conversely, CD147 showed a trend toward increased expression in forskolin and SMN:GFP expressing SMA astrocyte co-culture SG samples compared to healthy control SG samples, particularly so within the male SMA patient-derived samples (Fig 6H). Together, these data demonstrate that in combination with SMN gene therapy, forskolin treatment can further enhance SMN protein levels and influence expression of SYN1 within the SMA synaptogliosome.

**Figure 6.**
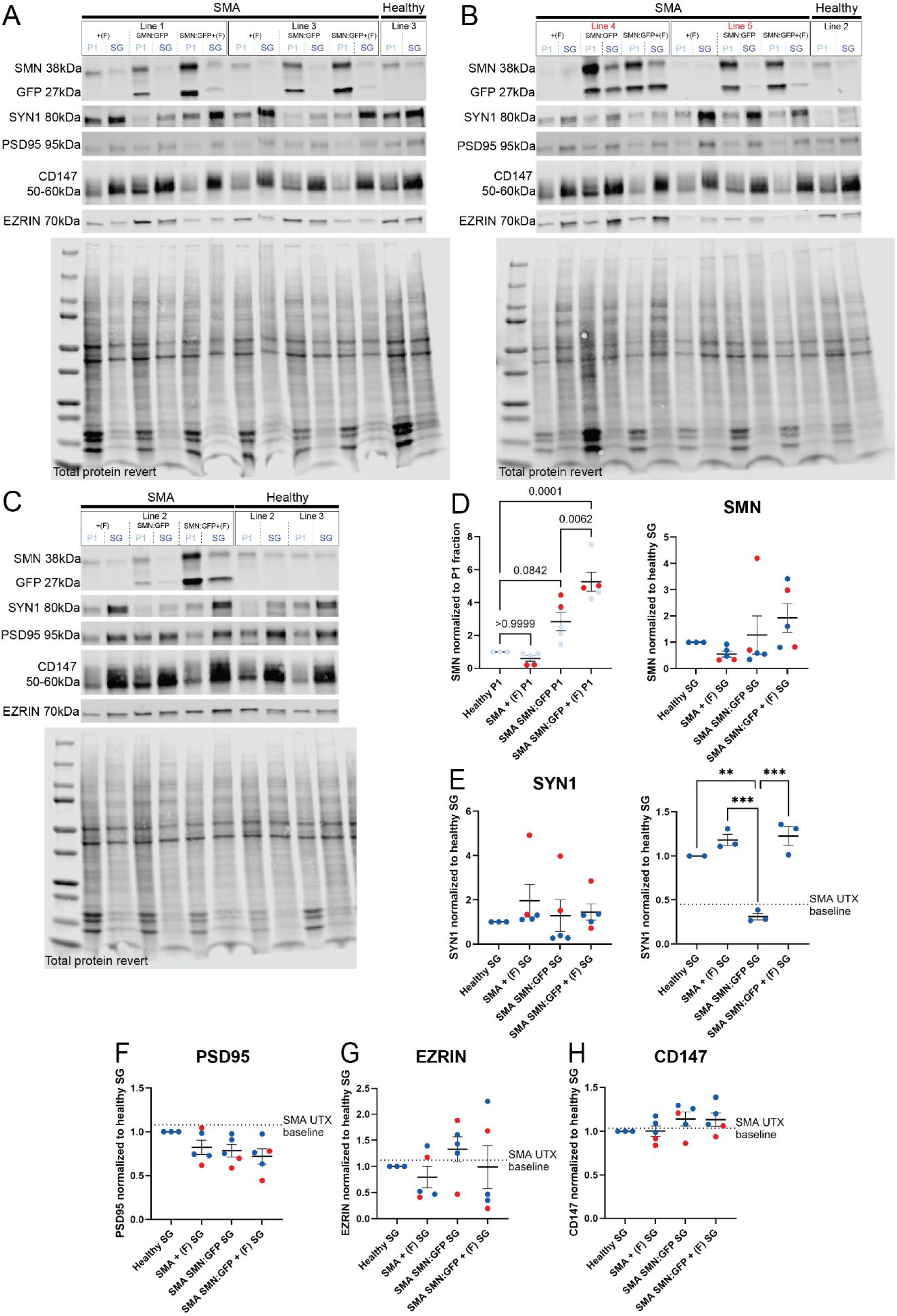
Forskolin treatment of SMA motor neuron astrocyte co-cultures restores synapsin I levels at the synaptogliosome. (A-C) Immunoblots of P1 and SG fractions from iPSC-derived healthy and SMA patient motor-neuron astrocyte co-cultures probed for SMN, synaptic (SYN1, PSD95) and PAP-associated (CD147, EZRIN) proteins. SMA co-cultures included the following conditions: baseline SMA cultures with forskolin (2-week duration), SMA cultures with SMN:GFP re-expressing astrocytes (with and without forskolin 2-week treatment). All protein signal was normalized to total protein revert signal. **(D)** Quantification and comparison of SMN protein levels in P1 fraction and SG fraction across all co-culture conditions. Quantification and comparison of SYN1 **(E)**, PSD95 **(F)**, EZRIN **(G)** and CD147 **(H)** protein levels in SG fractions across all co-culture conditions. One way ANOVA with Bonferroni multiple comparison statistical testing; **p<0.007; ***p=0.0001. N=8 (biological replicates), n= 1 (technical replicate).

We performed a similar analysis via immunocytochemistry to visualize synaptic protein alignment (SYN1/2; PSD95) and PAP-associated proteins (PEZRIN) in SMA co-cultures at baseline and with our dual treatment paradigm. Due to the stronger synaptic phenotype presentation in the male SMA patient lines, we utilized these lines in our downstream synapse/PAP quantification experiments. Within the co-cultures, we were readily able to detect EZRIN and pERM proteins specifically within astrocytes (Fig. 7A-D); EZRIN was observed within cytoplasmic regions and filopodia of astrocytes (Fig. 7A&C), while pERM proteins were highly enriched within astrocyte filopodia (Fig. 7B&D). This is consistent with previous findings demonstrating the selective enrichment of pEZRIN within astrocyte filopodia and PAP regions *in vivo* (8, 14). Immunocytochemistry analysis revealed the robust detection of pre- (SYN1/2) and post- (PSD95) synaptic terminal proteins and pERM proteins within healthy co-cultures (Fig. 7E), contrasting to the signal detected within SMA patient-derived co-cultures which remained sparse (Fig. 7F). We used the Imaris software to quantify synapses (total distance between SYN1/2 and PSD95 protein puncta ≤0.5µm) and predicted PAP regions across co-culture conditions (Fig. 7G&H). This revealed a significant reduction in number of synapses and predicted PAP region in SMA patient-derived co-cultures compared to healthy samples (Fig. 7I&J); this trend was observed across all other co-culture conditions with healthy/patient-derived cell combinations, suggesting both intrinsic defects in motor neurons and astrocytes contribute to this disease phenotype. Consistent with our synaptogliosome data, we found that combining SMN gene therapy with forskolin treatment significantly improved the detection of complete synapses (≤0.5µm distance between SYN1/2 and PSD95 signal) compared to baseline SMA co-cultures (Fig. 8A-C). pEZRIN protein expression showed mild improvements in the dual treatment paradigm SMA co-cultures, although we found that forskolin induced a significant decrease in pEZRIN protein in healthy co-cultures (Fig. 8A, B, D). Experiments completed with SMN:GFP expressing SMA astrocytes demonstrated that forskolin modulated astrocyte cell morphology within the co-culture system, with forskolin treated cultures demonstrating elongated astrocyte morphology with increased branching and fine processes (Fig. 8E&F).

**Figure 7.**
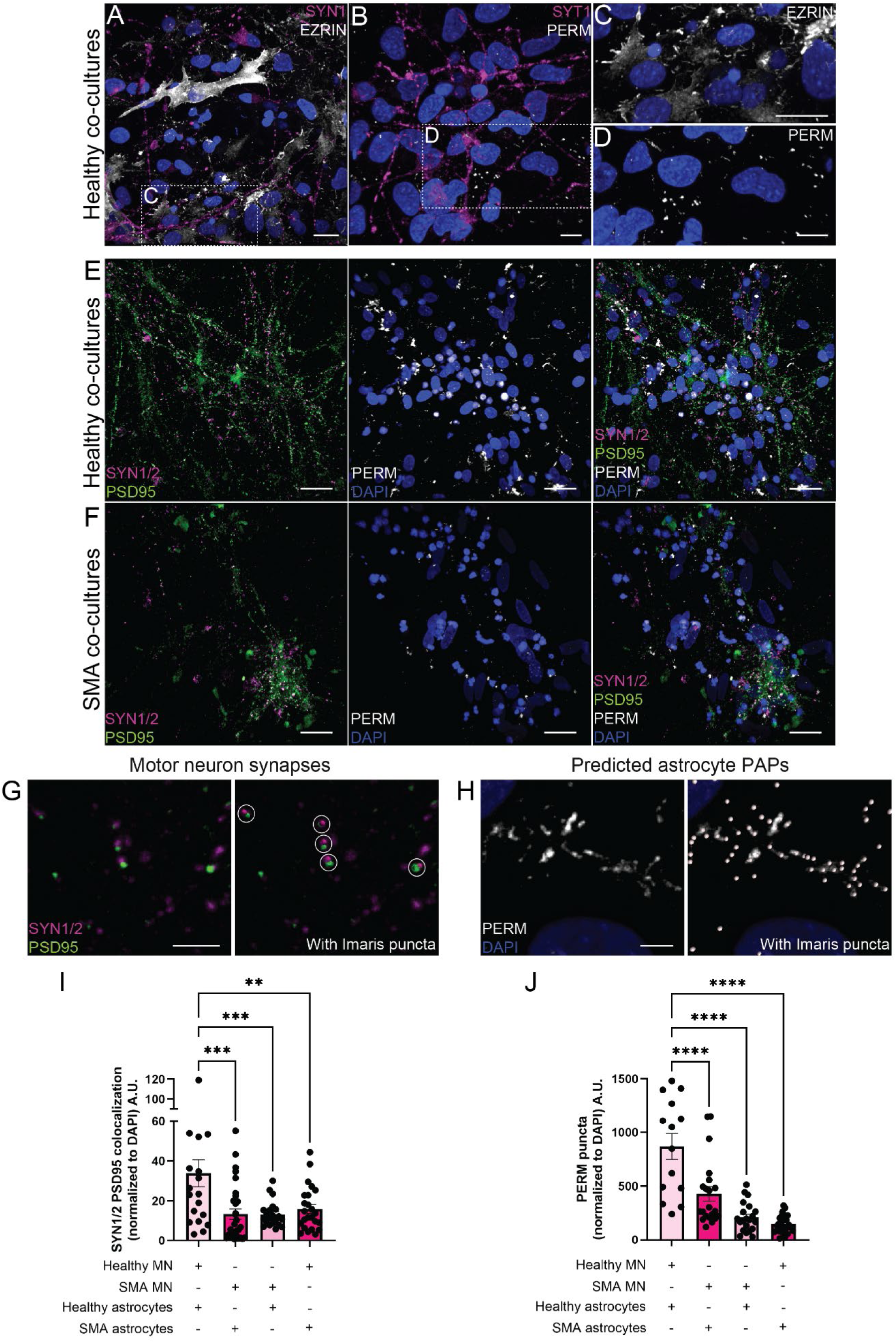
SMA patient-derived co-cultures demonstrates decreased number of synapses and PAP-associated proteins. Confocal images of healthy motor neuron-astrocyte co-cultures demonstrating enriched signal of **(A&C)** EZRIN and **(B&D)** phosphorylated ERM within astrocyte filopodia. Scale bars in A: 20µm, B: 10µm, C: 20µm, and D:10 µm. Confocal images of pre-(SYN1/2; magenta) and post- (PSD95; green) synaptic terminal proteins, and PAP-associated protein, PERM (grey signal) in **(E)** healthy and **(F)** SMA patient motor neuron astrocyte co-cultures. Scale bars: 30µm. Example of Imaris software puncta identification for **(G)** synaptic SYN1/2 PSD95 signal and **(H)** PAP-associated PERM signal. Quantification of **(I)** co-localized SYN1/2 PSD95 puncta (distance between puncta ≤0.5µm) and **(J)** PERM puncta in healthy and SMA patient motor neuron astrocyte co-cultures. Puncta number was normalized to DAPI signal (blue) observed per image. One-way ANOVA with Bonferroni multiple comparison testing; ****p<0.0001; ***p<0.0008, **p=0.0026. N=5 (biological replicates), n= 3 (technical replicates).

**Figure 8.**
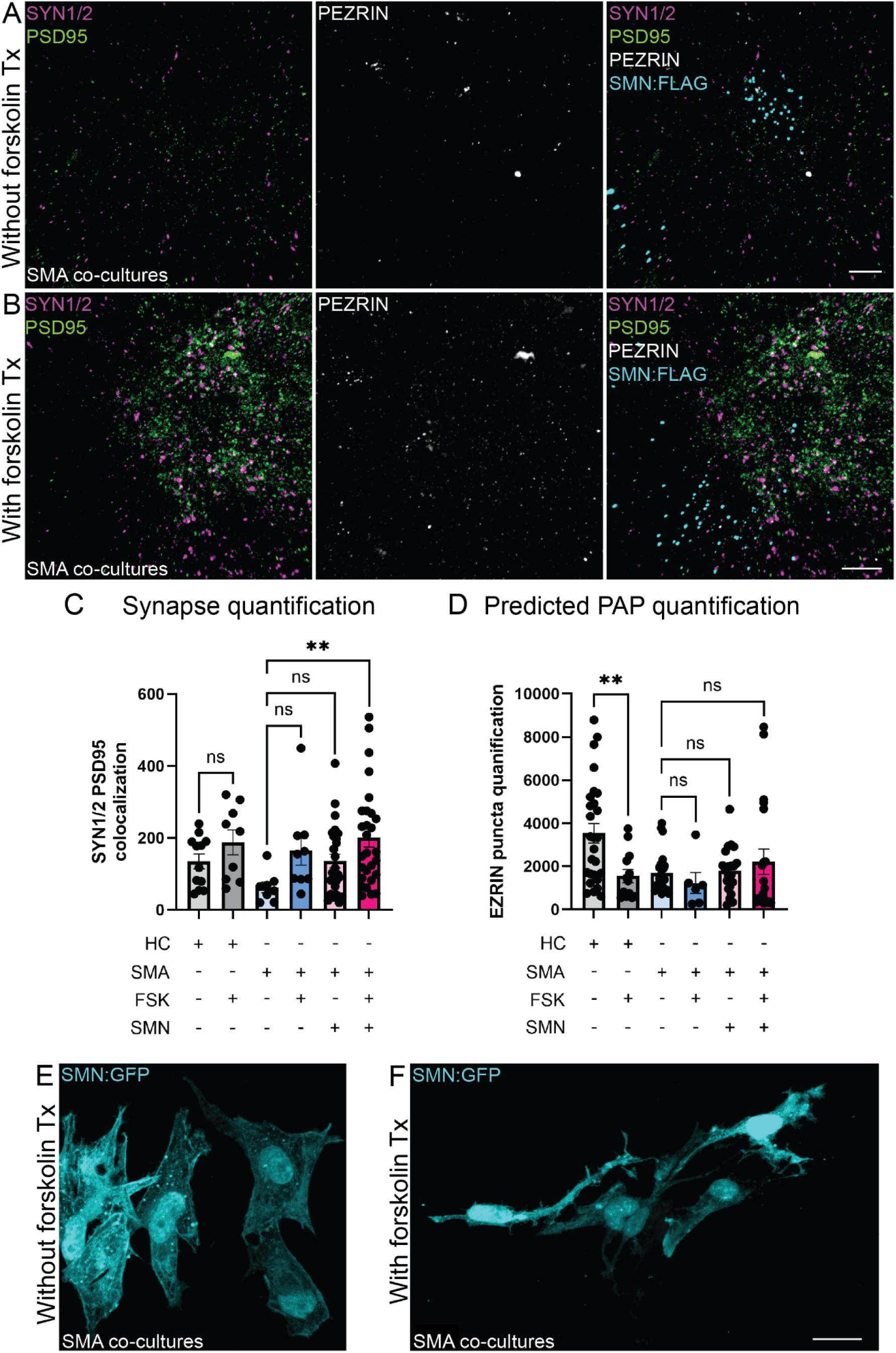
Forskolin treatment and astrocyte-targeted SMN gene therapy improves synapse quantification in SMA motor neuron co-cultures. Confocal images of pre- (SYN1/2; magenta) and post- (PSD95; green) synaptic terminal proteins, PERM PAP associated proteins (grey) and SMN:FLAG signal (cyan) in SMA motor neuron co-cultures containing SMN re-expressing astrocytes without forskolin treatment **(A)** and with forskolin treatment **(B)**. Scale bars in A&B: 10µm. Quantification of **(C)** co-localized SYN1/2 PSD95 puncta (distance between puncta ≤0.5µm) and **(D)** PERM puncta in healthy and SMA patient motor neuron astrocyte co-cultures. One-way ANOVA with Bonferroni multiple comparison testing; **p≤0.008. Confocal images of SMN signal (cyan) in SMA motor neuron co-cultures containing SMN re-expressing astrocytes (SMN:GFP) without forskolin treatment **(E)** and with forskolin treatment **(F)**. Scale bars in F: 15µm. N=5 (biological replicates), n= 3 (technical replicates).

Lastly, to assess if the dual treatment paradigm translated into improved function in SMA patient-derived motor neurons, we performed whole-cell voltage-clamp recordings and multi-electrode arrays to assess electrophysiology properties of motor neurons. We first examined passive membrane properties of motor neurons, including membrane capacitance, resting membrane potential (RMP), and input resistance (R_in_). Membrane capacitance did not differ significantly among the five groups (Fig. 9A), suggesting that motor neuron cell size and membrane surface area were not significantly altered by the different co-culture conditions. Under current-clamp conditions, RMP measured at 0 pA holding current was also similar across all groups (Fig. 9B). To assess membrane input resistance, hyperpolarizing current steps (-40 pA, 10-pA increments, 500 ms duration) were injected into recorded motor neurons (Fig. 9C). Plotting the peak membrane voltage responses against injected current revealed a significant increase in R_in_ in SMA motor neurons co-cultured with SMA astrocytes compared to healthy co-cultures. This elevation in R_in_ was reversed in SMA co-cultures via forskolin treatment or by SMN re-expression within SMA astrocytes, with or without additional forskolin treatment (Fig. 9C-E). We next examined whether these treatments affected neuronal excitability by measuring action potential firing in response to depolarizing current injections (Fig. 9F). SMA motor neurons co-cultured with SMA astrocytes exhibited a significantly higher action potential firing frequency compared to healthy co-cultures. This increase in firing of SMA motor neurons was attenuated by forskolin treatment and by co-culture with SMN re-expressing SMA astrocytes, either alone or in combination with forskolin (Fig. 9F&G). To further assess excitability, we measured the rheobase, defined as the minimal depolarizing current required to elicit an action potential. SMA motor neurons co-cultured with SMA astrocytes exhibited a reduced rheobase compared with the healthy control group (Fig. 9H). This reduction was rescued only by the combined SMN re-expression within SMA astrocytes and forskolin treatment; neither treatment alone was sufficient to normalize the rheobase in SMA astrocyte co-cultured motor neurons (Fig. 9H).

**Figure 9.**
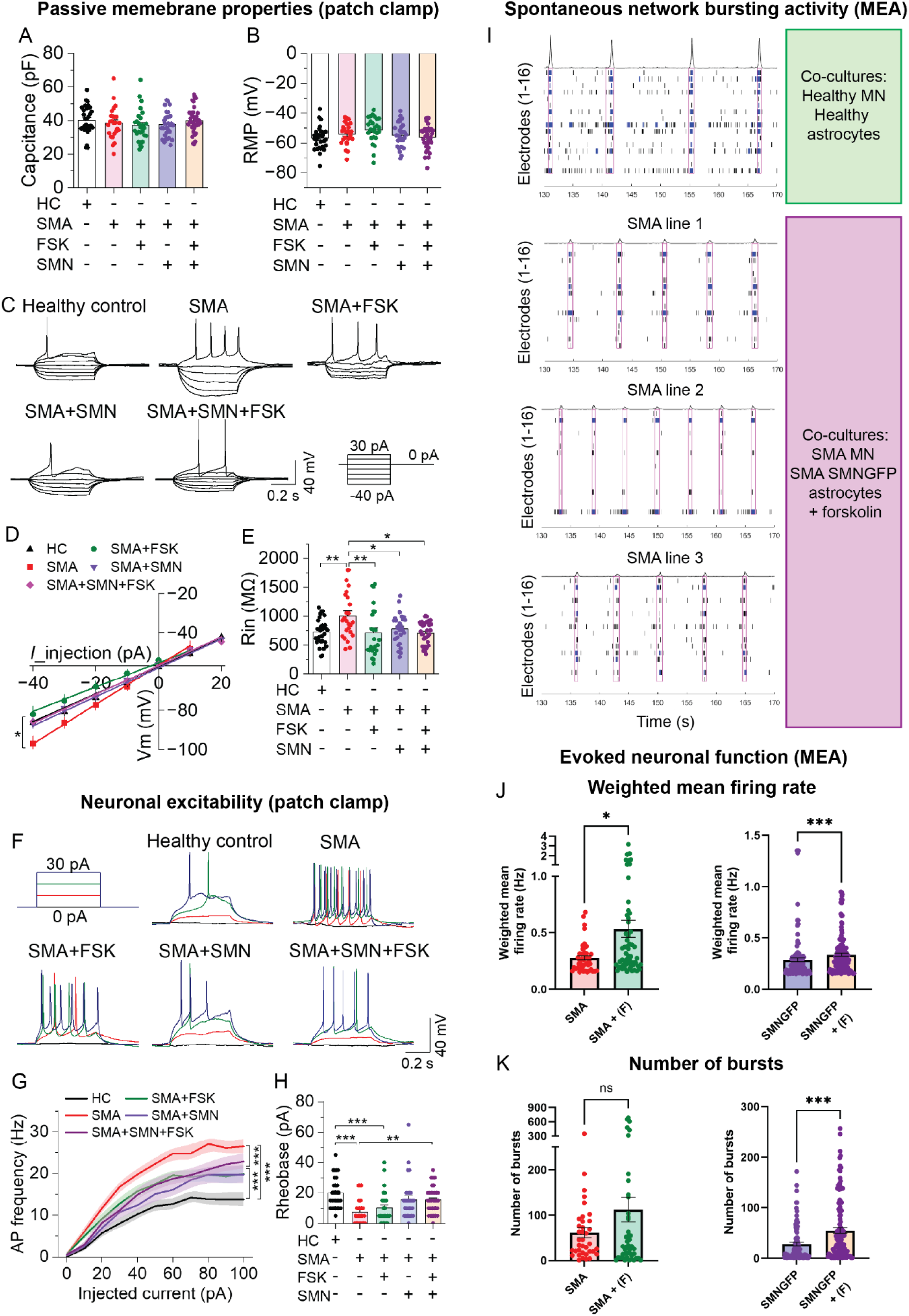
Improvements in SMA motor neurons function with forskolin treatment and SMN re-expressing SMA astrocytes. Membrane capacitance **(A)** and resting membrane potential (RMP) measurements **(B)** across co-culture groups (HC: healthy cultures, SMA: patient cultures; FSK: 10uM forskolin treatment for 2-week duration, SMN: culture with SMN re-expressing SMA astrocytes). **(C)** Representative membrane voltage traces in response to step current injections. Current–voltage relationship of steady-state membrane potential responses **(D)** and summary of input resistance (R_in_) **(E)** showing motor neurons co-cultured with SMA astrocytes exhibited significantly higher R_in_ compared with healthy co-culture group. This increase was reversed by treatment of forskolin (FSK) and SMN:GFP lentivirus transduction, with or without additional FSK treatment (One-way ANOVA; \**p*<0.05, \*\**p*<0.01). **(F)** Representative action potential (AP) traces in response to step current injections. **(G)** Plot of AP frequency in response to injected current across groups. SMA motor neurons co-cultured with SMA astrocytes exhibited a higher frequency of action potential firing in response to current injections compared to healthy cultures. This increase was reduced by FSK treatment and by SMN:GFP lentivirus transduction, with or without additional FSK treatment (Two-way repeated measures ANOVA; ***p < 0.001). **(H)** SMA astrocyte co-cultured SMA motor neurons, with or without FSK, exhibited decreased rheobase, and this decrease was rescued by co-culture with SMN:GFP lentivirus with FSK treatment (One-way ANOVA; \*\**p*<0.01, \*\*\**p*<0.001). **(I)** Rater plots of network bursting activity from spontaneous multi-electrode array recordings for healthy co-cultures, and SMA motor neuron co-cultures containing SMN re-expressing SMA astrocytes with forskolin treatment. Quantification of weighted mean firing rate **(J)** and number of bursts **(K)** from electrically evoked multi-electrode array recordings across all co-culture conditions. Mann-Whitney test; *p=0.0146, *** p=0.0002. N=5 (biological replicates), n= 2 (technical replicates).

We next assessed SMA motor neuron synaptic connectivity via multielectrode array across the different treatment paradigm conditions. Spontaneous network bursting events, indicating synchronized motor neuron synaptic communication across multiple electrodes, were readily detected within healthy co-cultures and within SMA motor neuron co-cultures containing SMN re-expressing SMA astrocytes and forskolin treatment (Fig. 9I). The other SMA motor neuron co-culture conditions (baseline, forskolin treatment alone, SMN re-expressing SMA astrocytes) did not demonstrate spontaneous network bursting activity (data not shown). We previously demonstrated that neither weighted mean firing rate nor bursting events of healthy motor neurons was improved by co-culture with SMN re-expressing SMA astrocytes (35); hence, we focused on assessing the effects of forskolin treatment within the SMA co-cultures of this current study. Forskolin treatment led to a significant increase in weighted mean firing rate of SMA motor neuron co-cultured withSMA astrocytes or SMN re-expressing SMA astrocytes (Fig. 9J). While forskolin led to an increasing trend in the number of bursting events in SMA motor neuron co-cultures with SMA astrocytes, which is indicative of action potential organization at individual electrodes, a significant increase in bursting events was only apparent in forskolin and SMN re-expressing co-treatment conditions (Fig. 9K). Together, our combined electrophysiological data highlight that a dual therapeutic approach consistently allows for significant improvements in intrinsic properties and network communication of SMA motor neurons.

## Discussion

Mechanisms underpinning tripartite synapse dysfunction at the central afferent synapses involved in SMA remains to be fully elucidated. Here, we defined novel intrinsic actin defects within SMA astrocytes that impact expression of pERM and EAAT1 proteins within filopodia and peri-synaptic astrocyte processes. We also molecularly dissect the synaptogliosome from SMA co-cultures to reveal a specific downregulation of synaptic vesicle release protein, SYN1, within the pre-synaptic terminal of motor neurons, which is associated with a significant decrease in synapse formation and synaptic dysfunction. Moreover, we found that combining astrocyte targeted SMN re-expression and forskolin treatment, utilizing therapeutic approach to target SMN-dependent and independent/related disease mechanisms, improves motor neuron synaptic protein expression and electrophysiological function. Further work is required to investigate restorative therapeutic strategies to correct PAP defects in SMA.

### Non-cell autonomous disease mechanisms: intrinsic astrocyte defects related to actin remodeling in SMA

Previous studies have mostly studied actin dynamics within *in vitro/in vivo* motor neurons (40–42) and neuronal cellular models (e.g., PC12 cells (43, 45), NSC34 cells (44)) to demonstrate that SMN deficiency leads to defective axonal growth, neurite outgrowth and growth cone collapse and downstream disruption of actin signaling pathways. We provide additional evidence that actin dynamics are perturbed in human SMA patient-derived astrocytes, which primarily influences the levels of actin regulating proteins (e.g., TENM2, TENM4, THY-1, CD44, NRCAM) at the plasma membrane required for instigating filopodia motility. The reduced level of these proteins likely underlies the failure of the SMA patient-derived astrocytes to dynamically respond to acute actin remodeling stimulation and modulate their filopodia density and levels of active CDC42-GTP, the small RhoA GTPases that primarily regulates filopodia actin dynamics. This finding is consistent with a previous study demonstrating lower levels of active CDC42-GTP with SMN deficiency (43).

Other RhoA GTPases may contribute to this phenotype in SMA astrocytes, since SMN deficiency also correlates with increased RhoA levels (43), which can further alter astrocyte morphology by inducing filopodia withdrawal and decreasing finer astrocyte processes (63). Our previous work (49) and the observation that several cell surface proteins associated with glial inflammatory responses are significantly increased in abundance in SMA astrocytes (Fig. 1), demonstrates that SMA patient-derived astrocytes are reactive, which can further influence cell morphology and process dynamics even within an *in vitro* setting (64). We noted the more typical polygonal and less stellated morphology across our *in vitro* astrocyte cultures; however, at baseline SMA astrocytes appeared swollen with prominent F-actin stress fibers compared to healthy astrocytes, suggesting global alterations in actin cytoskeletal dynamics in SMN deficiency. This is consistent with a previous study demonstrating SMN deficiency is associated with increased cell area, rounded cell morphology and F-actin fraction (44).

Based on our proteomics analysis, we hypothesized that the filopodia actin-associated phenotypes likely extend to PAPs, which tightly regulates neuronal function and synaptic plasticity via direct contact interactions. This represented an intriguing avenue for further exploration, since we previously demonstrated SMN restoration alone was not fully sufficient to restore astrocyte intrinsic defects driving diminished motor neuron activity (35). We decided to further explore the disease mechanism involving CD44, due to the previous associations with excitatory synapse maturation and refinement, and regulation of astrocyte morphology via regulation of RhoA GTPases signaling (22, 65), and physical interactions with PAP-associated protein, pERM, (17–19) at the plasma membrane. Previous reports indicate high expression of CD44 in reactive astrocytes in neurodegenerative diseases affecting different brain regions, including epilepsy-related disorders (66), Alzheimer’s disease (67) and prion disease (68), while we observed a significant decrease in surface-localized CD44 abundance in spinal cord patterned SMA astrocytes compared to healthy samples. While decreasing CD44 levels acts to reduce astrocyte reactivity and improves disease phenotypes in affected brain regions (66), it also was noted to decrease astrocyte-synapse contacts, indicating the potential role of CD44 in determining astrocyte coverage of synapses (69). With the reports of astrocyte CD44+ processes in close contact with anterior horn motor neurons of the spinal cord (70) and its previous defined roles in synaptic refinement (22, 65), it is possible that lower levels of CD44 in SMN deficiency acts to disrupt spinal astrocyte-motor neuron interactions. Hyperphosphorylated ERM phenotypes have also been reported to decrease neurite outgrowth (71), and reduce complexity of astrocyte morphology and excitatory synapse function in Parkinson’s disease (27), which can be alleviate by inhibiting phosphorylation of ERM proteins. However, spinal cord tissue derived from human ALS patient and rodent models demonstrates selective loss of tripartite synapses, with significant decreases in the levels of PAP-associated proteins, pEZRIN and glutamate transporter EAAT2 (25). These findings are more consistent with our human spinal cord patterned motor neuron astrocyte co-culture model of SMA, where we also demonstrate low expression of pEZRIN and EAAT1 within astrocyte filopodia regions. In relation to other molecular mediators of PAP-mediated regulation at the synapse, we identified other cell surface *N*-glycoproteins with previous associations with synapse formation and integrity (e.g., TNC, NECTRIN3 (72), PCDH17(73)) that showed lower abundance in SMA astrocytes. This also included ephrin receptors (e.g., EPHA3), which has previously been implicated in SMA astrocyte-mediated synaptic dysfunction in SMA (74). Applying newer capture methods and proximity labelling approaches (75, 76) to motor neuron-astrocyte co-cultures would provide further insight into protein-protein interactions at the synaptogliosome and how they are defective in SMA.

### Synaptogliosome protein perturbation in SMA patient-derived samples: consequences for regulation of motor neuron function

We have previously demonstrated that motor neuron activity is severely diminished in co-cultures featuring SMA-patient derived astrocytes which is exacerbated by reduced EAAT1 expression (35). At the single cell level, here we demonstrate via patch clamp electrophysiology that motor neurons within SMA astrocyte co-cultures are hyperexcitable, demonstrating increased input resistance, higher action potential frequency and reduced rheobase, which is a phenotype consistent with previous assessment of human SMA motor neurons (77). This is concomitant with a significant reduction in SYN1 expression, a severe reduction in motor neuron synapses, and decreased levels of PAP proteins associated with regulating neurotransmission (e.g., EAAT1), motility to respond to synaptic needs (e.g., pERM), and synapse formation (i.e., EPHA3, NECTRIN3). Our working hypothesis is that SMA motor neurons demonstrate impaired synapse formation and/or synaptic loss which likely is exacerbated by abnormal PAP dynamics of SMA astrocytes. Of the synapses that do form, there is also likely impaired glutamate neurotransmission, contributed to by reduced SYN1 levels in the pre-synaptic terminal and EAAT1 within astrocytic PAPs, which acts to induce an abnormal motor neuron hyperexcitable phenotype as part of a compensatory mechanism previously described in other neurodegenerative disease (78). When assessing function network maturation over time, motor neuron synaptic connectivity is subsequently impaired potentially due to excitotoxicity and/or failure of synapses to be formed and maintained properly. A caveat of our current system is that tripartite synapses are likely formed of motor neuron pre- and post-terminals, whereas sensory-motor circuitry within central afferent synapses would be formed of pre-synaptic proprioceptive neurons/interneurons and post-synaptic motor neurons *in vivo*. However, previous *in vivo* reports assessing proprioceptive innervation on to lower spinal motor neuron demonstrate a similar phenomenon whereby the reduction of the pre-synaptic neurotransmission from the sensory input can drive opposing effects on motor neuron excitability by increasing input resistance and reducing spiking ability (79).

Pre-synaptic defects involving synaptic vesicles have been consistently described in SMA disease phenotypes, affecting proprioceptive neurons (79, 80), interneurons (81), and motor neurons (82–85). Intriguingly, although it is a small sample size, we found that the SYN1 phenotype was more severe in male-derived vs female-derived cultures. Although the sex differences in SMA disease pathology remains to be fully elucidated, there is some evidence to suggest male patient present with a more severe disease phenotype (86, 87) especially since many of the SMA disease modifier genes, such as *PLS3*, *USP9X* and *UBA1*, are encoded on the X chromosome. We also note that protein detection at the cell surface in our proteomics approach maybe be influenced by sex differences (e.g., comparison between female healthy samples vs male SMA patient samples) in addition to SMN deficiency. Our observations would require further validation in future studies by increasing the number of SMA patient iPSC lines and screening all female lines for expression levels of known X-linked SMA disease modifiers.

Although we observed a significant reduction of pERM detection within the SMA co-cultures and EAAT1 in SMA astrocytes, other defined PAP-associated proteins within the synaptogliosome fraction, including CD147 and EZRIN, did not show significant differences between healthy and SMA samples. These data suggest that PAPs are still present within SMA conditions, but they may have specific intrinsic defects influencing certain protein levels and their downstream pathways. This phenomenon has been determined for mRNAs that undergo local translation within in PAPs of hippocampal astrocytes during learning and memory processing, whereby specific mRNAs content within PAPs changes after fear conditioning (9). Within our proteomics data, we noted the absence of protein detection within a significant proportion of SMA samples for specific low abundant cell surface proteins (e.g., TENM2, TENM4). Together with the previous reports of perturbated mRNA translation occurring with SMN deficiency (40, 56, 88–90), we hypothesize that there are likely local translation defects occurring within SMA PAPs that may underlie the lower abundance of specific cell surface proteins. Other have demonstrated that the F-actin bundling protein and SMN disease modifier PLS3 can improve the cell surface translocation of tropomyosin receptor kinase B, leading to enhanced TrkB/BDNF signaling and improved F-actin dynamics within the axon terminal of SMA motor neurons (91), demonstrating a critical link between cell surface proteins, actin dynamics, and SMN levels.

### Novel therapeutic strategy to restore human SMA motor neuron function

We demonstrated that dual application of forskolin treatment and astrocyte-targeted SMN1 gene augmentation led to the most significant improvements across the astrocyte intrinsic and co-culture phenotypes observed in this study. Forskolin, an adenylyl cyclase activator leading to increased cyclic AMP, was chosen due to its previous associations with stimulating CDC42-associated actin remodeling via protein kinase A pathway (59, 60) and enhancing astrocyte stellation via cAMP elevation (92, 93). Our rationale was that improved actin dynamics within SMA astrocytes by simultaneously targeting SMN-mediated disease mechanisms and actin dynamics would lead to improved astrocyte branching and filopodia density and better interaction of PAPs with motor neurons.

Both forskolin treatment and SMN1 gene replacement independently demonstrated improvements in SMA astrocyte filopodia density after actin stimulation, although for both conditions astrocyte cell morphology remained largely polygonal with swollen cell soma indicating partial phenotype restoration. Only SMN re-expressing SMA astrocytes with forskolin treatment demonstrated full stellation of SMA astrocytes with an increased number of filopodia, indicating improvements in actin dynamics to instigate full morphological stellation. This is consistent with previous observations that SMN restoration only partially rescues SMA astrocyte phenotypes (35, 57), and that additional therapeutic interventions likely are required for further phenotypic improvement (94).

By using a multi-pronged approach to probe the SMA synaptogliosome, we observed the different aspects of therapeutic benefits from applying a dual therapeutic paradigm. Even without SMN re-expressing astrocytes, forskolin treatment alone within SMA co-cultures dramatically improved SYN1 protein levels at the pre-synaptic terminal, synapse number, and individual motor neuron cell function by restoring input resistance and action potential frequency. Forskolin treated SMA co-cultures without SMN gene therapy also did not show increased SMN protein levels compared to untreated SMA cultures. This suggests a mechanism where forskolin can act independently to restore neurotransmitter release at the pre-synaptic terminal in a motor neuron cell autonomous manner, potentially via correcting actin dynamics. Similar phenotypes have been described related to abnormal F-actin and synaptic vesicle dynamics, such as docking, density and release via exocytosis, at the pre-synaptic terminal of neurons in SMN deficiency (82, 85, 95, 96), which can be modulated via influencing expression of the actin-binding protein, profilin2a (41, 45, 97). With cAMP being an important second messenger involved in many downstream signal transduction pathways, it is highly possible that other cascades (e.g., gene expression via CREB activation, MEK/ERK signaling, PI3K/Akt signaling) contribute to the phenotype restoration in both SMA motor neurons and astrocytes.

Interestingly, SMN protein levels did increase in co-cultures containing SMN re-expressing astrocytes with forskolin treatment, which suggests a synergistic effect of prolonged forskolin treatment on SMN levels and potentially greater SMN protein stability within astrocytes of these conditions (61). It is with these dual treatment conditions that we observe consistent improvements in motor neuron synapse number and electrophysiological function, leading us to hypothesize that forskolin may preferentially improves motor neuron pre-synaptic defects, while SMN re-expression provides partial restoration of astrocytic neuromodulatory functions to improve synaptic events. We previously demonstrated that restoring SMN levels in SMA astrocytes partially improves EAAT1 protein levels and glutamate uptake function (35), however in this study we demonstrate that SMN and GFP levels do not significantly increase within the synaptogliosome fraction. This suggests that other SMN-independent disease mechanisms could be involved in exacerbating phenotypes associated with the synapse and PAPs. Our data suggest that while forskolin promotes cell morphology changes and stellation in SMA astrocytes, other actin modulators may provide better outcomes for SMA PAP function. Other therapeutic strategies targeted to other PAP features, such as mitochondria, calcium signaling or local mRNA translation, might also be required to fully restore full PAP function. In the case of SMA motor neurons, therapeutic targeting of mRNA transport and local translation of synaptic proteins significantly improves neurotransmitter release and function (83), which an approach that may also be required for SMA PAP improvements.

### Conclusions

Overall, we demonstrate that SMA patient-derived astrocytes exhibit impaired filopodia actin dynamics that disrupts specific PAP protein expression (pERM and EAAT1), which likely exacerbates motor neuron synaptic dysfunction. While astrocyte targeted SMN re-expression alone is insufficient to fully restore PAP properties and downstream neuronal function, the addition of forskolin which acts to increase intracellular cAMP levels and actin dynamics, led to a significant improvement in motor neuron synapse formation and function. Together, this highlights the importance in defining the cell-autonomous and non-cell-autonomous disease mechanisms, and SMN-dependent and independent pathways involved in human SMA synaptopathy to generate novel therapeutic strategies.

## List of abbreviations

SMA: spinal muscular atrophy
SMN: survival of motor neuron
PAP: peri-synaptic astrocyte process
ERM: ezrin, radixin, moesin
PERM: phosphorylated ezrin, radixin, moesin
cAMP: cyclic adenosine monophosphate

## Declarations

### Ethics approval and consent to participate

The use of iPSCs was approved by the Medical College of Wisconsin Institutional Review Board (PRO00025822), the Institutional Biosafety Committee (IBC20120742) and the Human Stem Cell Research Oversight Committee.

### Consent for publication

We have obtained a Biorender publication license.

### Availability of data and materials

Proteomics data generated from this study are available in Massive (massive.ucsd.edu; dataset identifier: MSV00010125; password for reviewers: gundrylab1) and will be made publicly available upon publication of this manuscript. All other datasets used and/or analyzed during the current study are available from the corresponding author on reasonable request.

### Competing interests

The authors declare that they have no competing interests.

### Funding

This work was supported by the Audrey Lewis Young Investigator Award funded by CureSMA (E.W.), CureSMA (A.D.E), and the National Institutes of Health awards R35HL155460 and R01HL134010 (R.L.G.).

### Authors’ contributions

E.W. Conception and design, collection and assembly of data, data analysis and interpretation, manuscript writing, financial support. X.L. collection and assembly of data, data analysis and interpretation. L.B.L. collection and assembly of data. M.W. collection and assembly of data. R.L.G. collection and assembly of data, data analysis and interpretation, and financial support. Q-S.L. collection and assembly of data, data analysis and interpretation A.D.E. Conception and design, data analysis and interpretation, final manuscript editing, financial support.

## Acknowledgements

The Oxford Instruments Center for Advanced Microscopy-Electron Microscopy Core (RRID:SCR_026315) at the Medical College of Wisconsin is an institutionally available research service unit managed on behalf of the Medical College by the Department of Cell Biology, Neurobiology and Anatomy. This core provided the N-SIM super resolution microscope, and Imaris software license and PC used in this study. The SMN:FLAG lentiviral DNA constructs used in this study were kindly provided by the laboratory of Dr. Xue Jun Li (University of Illinois-Chicago). We also acknowledge the Viral Vector Core Facility at the MCW/BRI for generating the lentiviruses used in this study, the Cardiovascular Center equipment core at MCW for use of the MEA system, and Dr. Elizabeth Sweeney’s lab for use of their Optima Max-TL Ultracentrifuge. We acknowledge Greenstone Biosciences Inc. for providing the human iPSC line(s): GSB-2945, GSB-2394, GSB-2009. Mass spectrometry was performed using instrumentation in the CardiOmics Program supported by the UNMC Center for Heart and Vascular Research. The graphical abstract and schematic in Fig. 4A and Fig. 5A were generated using Biorender.

## Notes

### Competing Interest Statement

The authors have declared no competing interest.

